# Meiotic DNA breaks activate a streamlined phospho-signaling response that largely avoids protein level changes

**DOI:** 10.1101/2022.02.24.481857

**Authors:** Funda M. Kar, Christine Vogel, Andreas Hochwagen

## Abstract

Meiotic cells introduce a large number of programmed DNA breaks into their genome to stimulate meiotic recombination and ensure controlled chromosome inheritance and fertility. An intricate checkpoint network involving key kinases and phosphatases coordinates the repair of these DNA breaks during meiosis, but the precise DNA break-dependent phosphorylation targets remain poorly understood. It is also unknown whether meiotic DNA breaks change gene expression akin to the canonical DNA-damage response. To address these questions, we analyzed the meiotic DNA break response in *Saccharomyces cerevisiae* using multiple systems-level approaches. We identified 332 DNA break-dependent phosphorylation sites, vastly expanding the number of known DNA break-dependent phosphorylation events during meiotic prophase. Only about half of these events occurred in recognition motifs for the known meiotic checkpoint kinases Mec1 (ATR), Tel1 (ATM) and Mek1 (CHK2), suggesting that additional kinases contribute to the meiotic DNA break response. Surprisingly, the numerous changes in phosphorylation were accompanied by very few changes in protein levels despite a clearly detectable transcriptional response. To explain this dichotomy, we show that meiotic entry lowers the expression baseline of many mRNAs enough so that subsequent break-dependent mRNA production has no measurable effects on the largely stable proteome.

## Introduction

Homologous recombination during meiosis is initiated by programmed DNA breaks created by the conserved topoisomerase-like protein Spo11 (Bergerat et al., 1997; Keeney et al., 1997). By stimulating crossover formation, these DNA breaks promote genetic diversity in the offspring and, in many organisms, are essential for faithful segregation of chromosomes (Lam & Keeney, 2015). Accordingly, failure to produce DNA breaks leads to infertility in yeast and mammals (Baudat et al., 2000; Klapholz et al., 1985; Romanienko & Camerini-Otero, 2000). However, the large number of DNA breaks also represents an inherent hazard for genome instability. To protect the genome, a complex signaling network coordinates meiotic processes in response to DNA break formation and prevents inappropriate meiotic progression when break repair is delayed or defective (MacQueen & Hochwagen, 2011; Subramanian & Hochwagen, 2014).

We are only beginning to understand the signaling pathways that connect meiotic DNA break formation to DNA repair and other meiotic processes. Available data indicate a prominent role for the canonical DNA-damage sensor kinases ATR and ATM, although the extent to which the two kinases are linked to the control of meiotic processes may vary between organisms (Kar & Hochwagen, 2021). In the budding yeast, *Saccharomyces cerevisiae*, the homologues of ATR and ATM, Mec1 and Tel1, respectively, and the downstream CHK2-like effector kinase Mek1 regulate a large number of meiotic processes, including DNA break formation and repair, chromosome pairing, and meiotic cell-cycle progression (Kar & Hochwagen, 2021). Targeted studies in yeast have identified relevant DNA break-dependent phosphorylation events for several of these processes, including control of break levels (Carballo et al., 2013), suppression of sister-chromatid recombination (Callender et al., 2016; Carballo et al., 2008; Niu et al., 2009), crossover formation (Chen et al., 2015; He et al., 2020; Woo et al., 2020), centromere uncoupling (Falk et al., 2010), and control of meiotic cell-cycle progression (Chen et al., 2018; Penedos et al., 2015). Targeted mutagenesis based on known kinase motifs mapped additional, functionally important break-dependent phosphorylation sites (Bartrand et al., 2006; Cartagena-Lirola et al., 2006; Serrentino et al., 2013), and a number of DNA break-dependent, site-specific phosphorylation events of unknown functional significance have been identified (Kniewel et al., 2017; Shroff et al., 2004; Suhandynata et al., 2016). However, available data (Garcia et al., 2015; MacQueen & Roeder, 2009; Mohibullah & Keeney, 2017) indicates that our understanding of the signaling response to meiotic DNA break formation is far from complete.

One little-explored aspect of the meiotic DNA break response is the relationship between the signaling pathways discussed above and changes in gene expression. The canonical DNA-damage response in vegetative yeast cells involves a well-defined transcriptional response that is signaled through Mec1/Tel1-dependent activation of the effector kinases Rad53 and Dun1 (Jaehnig et al., 2013). These kinases activate a core set of DNA-damage response genes, including genes coding for DNA-repair factors and regulators of nucleotide abundance. Rad53 activity is attenuated, but not absent, during meiotic DNA break formation (Cartagena-Lirola et al., 2008), posing the question as to the role of transcriptome and proteome remodeling during this step in meiosis.

To address these questions, we conducted a systems-level analysis to capture the cellular response to meiotic DNA breaks with respect to phospho-proteomic, proteomic, and transcriptomic changes. We identified hundreds of novel DNA break-dependent phosphorylation sites, highlighting the breadth of the signaling response to meiotic DNA breaks. Surprisingly, we observed a transcriptional response to DNA breaks that did not lead to detectable changes in the proteome. We explain this discrepancy by the differences in regulation of protein and mRNA abundance during meiotic prophase.

## Results

### Experimental setup to measure multiple aspects of the meiotic DNA break response

To capture the meiotic DNA break response from multiple angles, we performed transcriptomic, proteomic, and phospho-proteomic analyses in *S. cerevisiae* (**Figure 1A**). We compared DNA break-proficient cells carrying a functional *SPO11* gene with catalytic *spo11-YF* mutants, which cannot form DNA breaks (Bergerat et al., 1997). To ensure the most robust and stringent analysis possible, we implemented several additional experimental features. First, to avoid differences in meiotic state between DNA break-competent *SPO11* cells, which delay in prophase compared to *spo11* mutants (Keeney, 2001), we removed the mid-meiosis transcription factor Ndt80 genetically from both strains. *NDT80* deletion synchronizes both cultures prior to the exit from meiotic prophase (Xu et al., 1995), and thus eliminates cell-cycle differences, which create a well-known false-positive signal when studying the cellular response to DNA damage (Gasch et al., 2001; Suhandynata et al., 2016). Second, to increase recovery of DNA break-dependent phosphorylation events, we used a *pph3Δ* mutation to inactivate protein phosphatase 4 (PP4), one of the major phosphatases responsible for erasing Mec1/Tel1- dependent phosphorylation marks (Falk et al., 2010; Hustedt et al., 2015; Keogh et al., 2006). Finally, to ensure reproducibility, proteomics and phospho-proteomics samples were collected from three independent biological replicates and analyzed by two complementary mass spectrometry techniques to maximize recovery of phosphosites. We used flow cytometric analysis of DNA replication as a proxy for meiotic synchronization to show that meiotic cultures completed S phase with similar kinetics within each set of biological replicates (**Supplemental Figure 1**).

**Figure 1:**
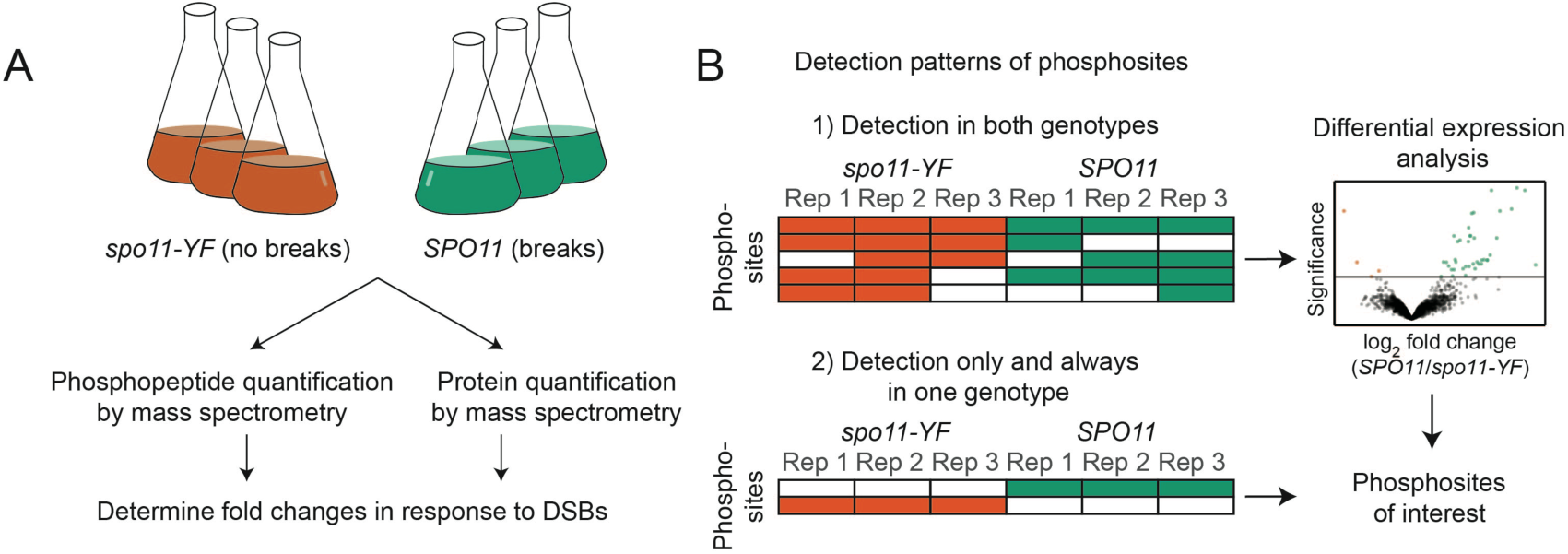
Workflow and experimental setup. (A) We extracted proteins from synchronous meiotic cultures and quantified the phosphopeptides and proteins obtained from *SPO11* and *spo11-YF* cells. Proteomics and phosphoproteomics experiments both had 3 biological replicates. (B) Quantitative and qualitative strategies were used to classify phosphorylation sites of interest. Filled rows represent non-zero intensity values for the phosphosites. When there were enough non-zero intensity values, differential expression analysis was used to classify DNA break-dependent sites or phosphorylation sites that were lost in response to DNA breaks (quantitative approach). In addition, we used a qualitative approach by analyzing detection patterns for phosphosites. When a phosphosite was present in all replicates of *SPO11* and absent in all *spo11-YF* samples, it was designated as a DNA break-dependent site. If a site was present in all replicates of *spo11-YF* and absent in all *SPO11* samples, it was categorized as lost in response to DNA breaks.

We quantified tryptic peptides before and after phospho-peptide enrichment to map DNA break-dependent changes in protein abundance and protein phosphorylation, respectively (**Figure 1A**). We used both Data-Dependent Acquisition (DDA) and Data-Independent Acquisition (DIA) to identify and quantify phosphorylation sites. DDA is the traditional method but is semi-stochastic as it only identifies the most abundant peptides. This filter biases the data towards high-abundance peptides and introduces variation in identified peptides between runs (Domon & Aebersold, 2010; Michalski et al., 2011). DIA overcomes this problem by co-fragmenting all of the peptides in pre-defined mass/charge windows. The resulting fragmentation spectra are highly complex and require more advanced algorithms and spectral libraries to resolve peptide sequences (Schubert et al., 2015), but DIA is able to achieve greater reproducibility and quantitative sensitivity than DDA (Bruderer et al., 2017; Selevsek et al., 2015), in particular for phosphoproteomics (Bekker-Jensen et al., 2020; Kitata et al., 2021). We designed a workflow that includes both methods to take advantage of their unique strengths.

### Meiotic DNA break response involves hundreds of phosphorylation events

We found high reproducibility across all three biological replicates for both the DDA and DIA proteomic analysis, and both methods efficiently recovered DNA break-dependent phosphorylation sites. We observed some loss of phosphorylation events in replicate 1, either during sample preparation or because of biological variability, as indicated by the narrow distribution phosphopeptide intensity differences between *SPO11* and *spo11-YF* cells in both the DDA and DIA analysis (**Supplemental Figure 2A,C**). Inspection of DNA content profiles showed that cultures from replicate 1 replicated their DNA slightly faster (**Supplemental Figure 1**), which may have resulted in longer prophase residence time, and led to loss of phosphorylation on some sites. This loss resulted in lower correlation of this replicate with the other replicates when examining all identified sites (**Supplemental Figure 2B,D**). However, the correlation with the other replicates was strong when considering only phospho-sites that were likely true-positive identifications, i.e. with an increased intensity in *SPO11* samples, suggesting this replicate produced meaningful data (**Supplemental Figure 3**). In addition to this, replicate 1 successfully reported previously characterized DNA break-dependent phosphorylation events like Zip1 S75 and Hed1 T40 (Callender et al., 2016; Falk et al., 2010), further confirming its validity. Therefore, we chose to retain the data from all three replicates. To ensure high-quality data, we removed phosphosite identifications with more than three values missing from the six measurements of DNA break-dependence.

We used two different approaches to establish the phospho-sites affected by DNA break formation (**Figure 1B**). First, we extracted phospho-sites detected in both *SPO11* and *spo11-YF* samples in at least three of the six samples and determined relative enrichment in *SPO11* versus *spo11-YF*. This analysis classified 40 and 157 phosphorylation sites as significantly enriched in the presence of meiotic DNA breaks (adjusted p-value cutoff 0.1) in the DDA and DIA data, respectively (**Figure 2A,B**). While these phosphorylation events are induced by DNA breaks, their presence in *spo11-YF* samples suggests that they can occur independently of meiotic DNA breaks as well.

**Figure 2:**
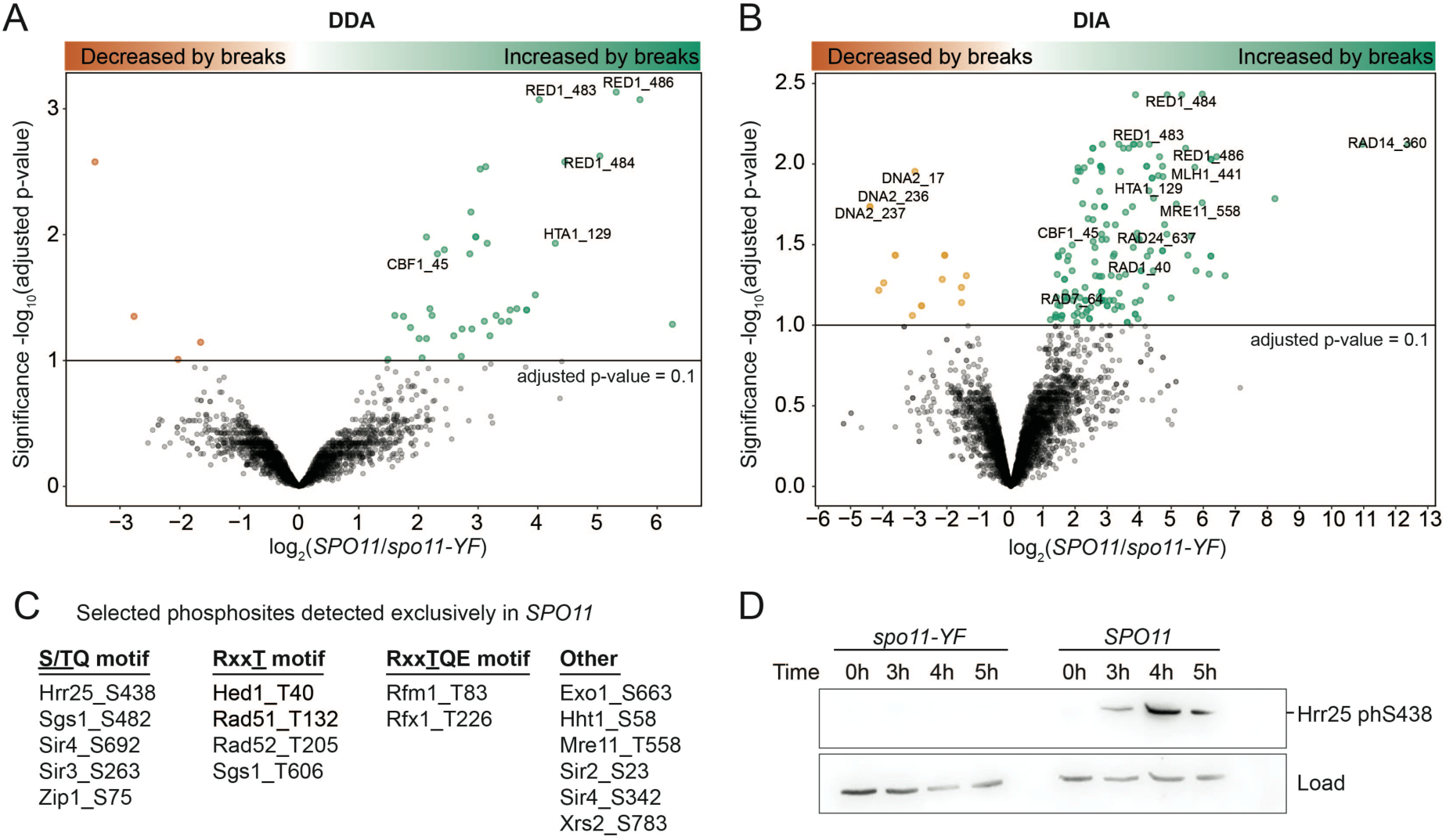
Changes in protein phosphorylation in response to meiotic DNA breaks. (A) A volcano plot showing results of differential expression analysis for DDA data, each dot representing a phosphosite. -log_10_ (Benjamini & Hochberg adjusted p-value) is shown on the y-axis and log_2_ fold change is shown on the x-axis. (B) A volcano plot showing results of differential expression analysis for DIA Data, each dot representing a phosphopeptide. Labels indicate the phosphorylation sites on peptides. (C) A table listing selected DNA break-dependent phosphorylation sites discovered by the presence/absence analysis; sites in black have been previously described (see Supplemental Table 3 for the complete list). (D) A western blot showing DNA break dependence of Hrr25 S438 phosphorylation. β-actin was used as a loading control.

Second, we conducted a presence/absence analysis to also include phosphosites that are never detected in *spo11-YF* samples, i.e. phosphosites that are likely fully dependent on meiotic DNA break formation (**Figure 1B**). This analysis revealed another 118 and 111 sites as phosphorylated only and always in presence of meiotic DNA breaks in the DDA and DIA data, respectively. In total, DDA and DIA analysis resulted in 158 and 241 DNA break-dependent phosphosites, respectively, with 67 sites captured by both methods.

We examined the method-specific site identifications and found that most were explained by missing data or scores below the significance cutoff (**Supplemental Figure 4**). Based on the results of these analyses, we opted to merge all DNA break-dependent phosphosites from the DDA and DIA datasets, obtaining a total of 332 DNA break-dependent phosphosites (**Supplemental table 3**). In addition, we detected 4,953 phosphorylation events that were not DNA break-dependent.

Among the DNA break-dependent sites were several previously confirmed Mec1/Tel1 targets, including H2A S129, Cbf1 S45, Zip1 S75, Rad54 T132, and Hed1 T40 (Callender et al., 2016; Downs et al., 2000; Falk et al., 2010; Niu et al., 2009; Smolka et al., 2007)(**Figure 2A-C**), confirming the high quality of our dataset. In addition, we uncovered many DNA break-dependent phosphorylation events that have only been characterized in non-meiotic cells or that are entirely novel. For example, we identified an additional DNA break-dependent phosphorylation event on histone H3 S57 (**Figure 2C**), which is predicted to weaken nucleosomal DNA association (Bowman & Poirier, 2015), and may thus play a role in removing nucleosomes during meiotic DNA repair. We also identified DNA break-dependent phosphorylation on several chromatin factors, including all three subunits of the Sir complex (Sir2, Sir3, Sir4), which have roles in meiotic checkpoint regulation and the timing of meiotic prophase (San-Segundo & Roeder, 1999; Subramanian et al., 2019)(**Figure 2C**). Among proteins involved in meiotic DNA repair, we identified additional DNA break-dependent phosphorylation sites on the MRX complex (Mre11, Xrs2), regulators of recombination (Rad52, Sgs1) and the meiotic resolvase complex (Mlh1, Exo1)(**Figure 2B,C**). Intriguingly, we also observed DNA break-dependent phosphorylation of multiple components of the nucleotide-excision repair machinery (Rad1, Rad7, Rad14, Rad16, Rad23, Rad26; **Figure 2B**). As there is no known role for nucleotide-excision repair during meiotic prophase, these phosphorylation events may be inhibitory.

We also observed a total of 81 phospho-sites that disappeared during DNA break formation, as they were either enriched in *spo11-YF* compared to *SPO11* samples or were specifically detected in *spo11-YF* samples (**Supplemental Table 4**). Amongst these sites were Cdk1-dependent phosphorylation events on Dna2 S17 and S237 (**Figure 2B**), which are known to regulate Dna2 localization to DNA breaks and may promote Mec1/Tel1-dependent phosphorylation of yet unidentified sites on Dna2 (Chen et al., 2011). Thus, the observed DNA break-dependent reduction in abundance of some of these phosphorylations is likely biologically meaningful.

Several known DNA break-dependent sites were absent from our data. Specifically, neither DDA nor DIA data captured Hop1 T318 or H3 T11 (Carballo et al., 2008; Kniewel et al., 2017). Inspection of the sequences surrounding these sites revealed that the H3 T11 phosphorylation site would reside on a 5 amino acid long tryptic peptide, which is too small for reliable proteomic identification. Conversely, a tryptic peptide with Hop1 T318 phosphorylation would be 49 amino acids long; and larger peptides are typically detected at a low frequency (Swaney et al., 2010).

Finally, we validated one newly identified site, namely the phosphorylation of the yeast casein kinase 1δ/ε homologue Hrr25 on S438 (**Figure 2C**) by raising and purifying a phospho-specific antibody (**Supplemental Figure 5**). Immunoblotting showed that Hrr25 S438 occurred specifically in response to meiotic DNA break formation and was undetectable in meiotic *spo11-YF* cultures (**Figure 2D**). Therefore, our data presents a high-quality, rich resource for phospho-signaling during the meiotic DNA break response.

### Phosphorylation sites are enriched for known kinase motifs

To better define substrate selection in response to meiotic DNA break formation, we examined features of the DNA break-dependent phospho-sites in our data. Motif enrichment analysis showed strong enrichment (motif-x, p-value cutoff 0.05) of predicted consensus sites for Mec1/Tel1 (<u>S/T</u>Q) and Mek1 (Rxx<u>T)</u>, the main regulators of the meiotic DNA break response. Approximately 19% of all DNA break-dependent sites occurred in an <u>S/T</u>Q motif, compared to only ∼5% <u>S/T</u>Q occurrence among all phosphorylation sites detected in our study (**Figure 3A**). Among DNA break-dependent <u>S/T</u>Q sites, L was the most common amino acid at the -1 position and E was the most common amino acid at the +2 position (**Figure 3B**). As L<u>S/T</u>QE is known as a high-affinity site for human ATM in *in vitro* studies (O’Neill et al., 2000), these sites are likely also targeted by the yeast orthologs Mec1/Tel1.

**Figure 3:**
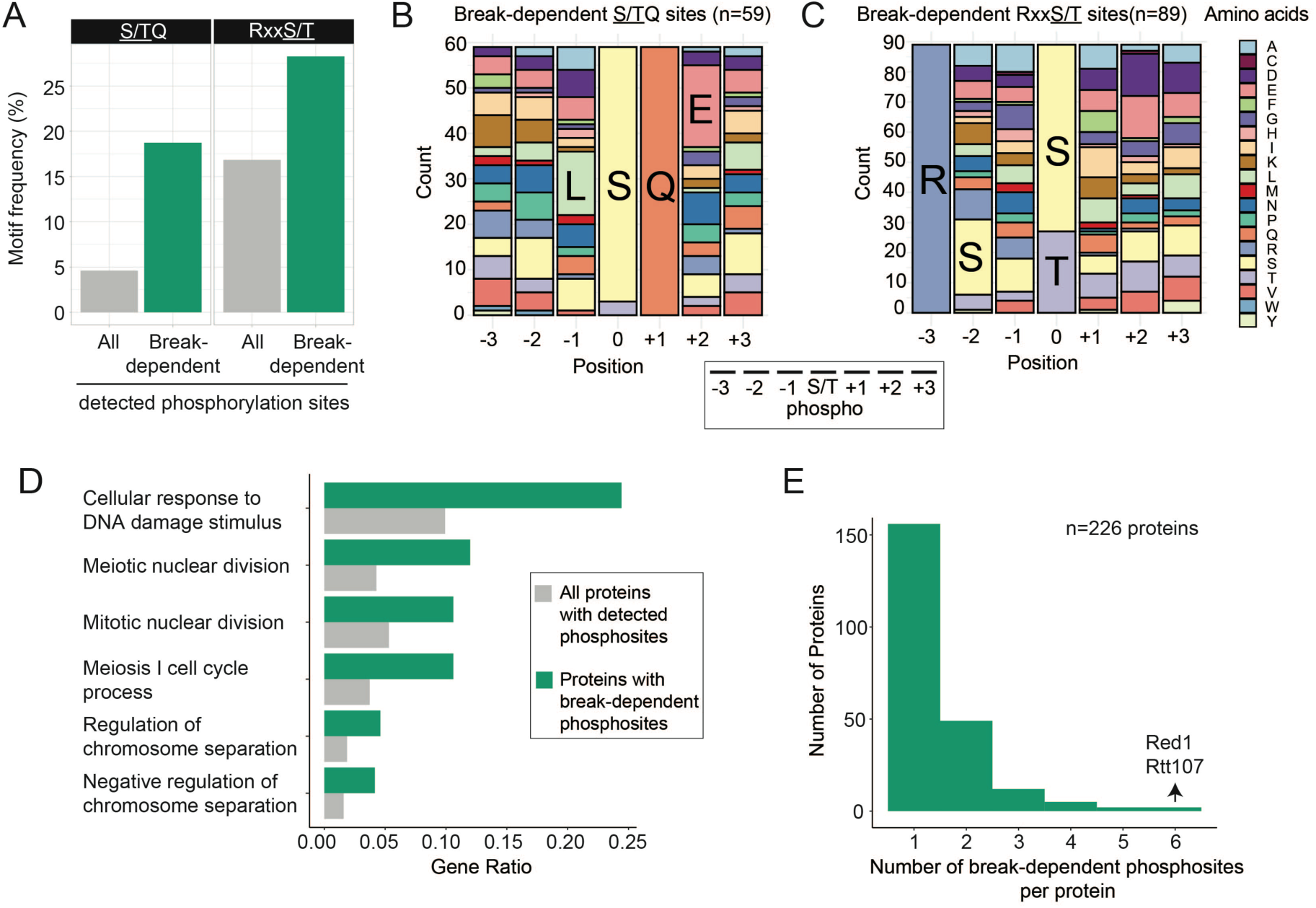
Characteristics of DNA break-dependent phosphorylation events. (A) DNA break-dependent sites were enriched for Mec1/Tel1 (<u>S/T</u>Q) and Mek1(Rxx<u>T</u>) consensus motifs, compared to all phosphorylation sites detected in our study. (B) Stacked bar graph representing the distribution of amino acids surrounding the DNA break-dependent <u>S/T</u>Q sites. The most frequent amino acids at the +1 and -1 position were leucine (L) and glutamic acid (E), respectively. (C) Stacked bar graph representing the distribution of amino acids surrounding the DNA break-dependent Rxx<u>S/T</u> sites. Serine (S) at the -2 position was the most frequent amino acid, while there was no striking feature for the other positions. (D) Bar-graphs showing the results of functional enrichment analysis of proteins with DNA break-dependent phosphosites (E) Bar graph showing the distribution of detected DNA break-dependent phosphorylation sites per protein.

Approximately 28% of all DNA break-dependent sites localized to Rxx<u>S/T</u> motifs, compared to only ∼17% of all detected sites (**Figure 3A**). Amongst these sites, we observed S as the most common amino acid at the -2 position (**Figure 3C**). The Rxx<u>S/T</u> motif is preferred by several kinases in yeast (Mok et al., 2010), but all known Mek1-dependent phosphorylation sites contain a threonine instead of a serine (Callender et al., 2016; Kniewel et al., 2017; Niu et al., 2009). Indeed, among the detected phosphorylation sites with an Rxx<u>S/T</u> motif, DNA break-dependent sites were significantly enriched for threonine as the phosphorylated amino acid (hypergeometric test, p-value <0.001; **Supplemental Figure 6A**), consistent with the reported sequence preference of Mek1 (Suhandynata et al., 2016).

In a couple of instances, we also observed phosphorylation of Rxx<u>T</u>QE sequences (**Figure 2C**), which combine Rxx<u>T</u> and <u>T</u>QE motifs and may thus represent a hybrid target motif for Mek1 and Mec1/Tel1. As Mek1 is activated by Mec1/Tel1, phosphorylation of Rxx<u>T</u>QE hybrid motifs by both types of kinases could create a coherent feedforward signal, a common feature in gene regulatory networks that creates signal stability (Mangan & Alon, 2003). Notably, the two proteins with phosphorylated RxxTQE motifs in our data, Rfm1 and Rfx1, are transcription factors of the DNA damage response, with Rfm1 preventing premature exit from meiotic prophase (McCord et al., 2003; Xie et al., 1999).

Although our data show a clear enrichment of predicted Mec1/Tel1 and Mek1 motifs, more than half (∼53%) of all DNA break-dependent phosphosites had neither motif, suggesting other kinases might be catalyzing these phosphorylation events. Motif search failed to identify additional significantly enriched motifs among these sites, possibly because of a lack of sequence preference or because of contributions from multiple kinases. Candidate kinases include Rad53 and Dbf4-dependent kinase (DDK), both of which have been shown to mediate phosphorylation events downstream of Mec1/Tel1/Mek1 and/or meiotic DNA break formation (Bashkirov et al., 2003; Chen et al., 2015; He et al., 2020; Lao et al., 2018), with only DDK having a known consensus motif.

In general, DNA break-dependent phosphorylation events localized to intrinsically unstructured parts of target proteins (**Supplemental Figure 6B**), presumably reflecting increased accessibility and the higher preponderance of serines and threonines in these regions. This trend is not unique to DNA break-dependent phosphorylation events (**Supplemental Figure 6B**) and has also been noted in previous large-scale phospho-proteomics analyses (Holt et al., 2009). Interestingly, almost one third (116 of 332) of DNA break-dependent phosphorylation events occurred within five amino acid of another DNA break-dependent site, which may either reflect multiple phosphorylation events mediated by the same kinase, or priming events, whereby phosphorylation by one kinase stimulates nearby phosphorylation events by another kinase.

The 332 phosphorylation sites mapped to a total of 226 different proteins which were strongly enriched for functions related to meiosis and DNA repair (q-value < 0.2, **Figure 3D**), implying that many of these phosphorylation events likely are functional. For most proteins, we only identified a single DNA break-dependent site, while two proteins had six DNA break-dependent phosphorylation sites (**Figure 3E**). One of these proteins is the meiotic chromosome organizer Red1, which has been suggested to be phosphorylated in both DNA break-dependent and -independent manner (Bailis & Roeder, 1998; de los Santos & Hollingsworth, 1999; Lai et al., 2011; Wan et al., 2004). Consistent with this notion, our data shows ten additional phosphorylation sites for Red1 that are not DNA break-dependent. One of the DNA break-dependent sites on Red1, T484, fits the consensus motif for phosphorylation by Mek1, but phosphorylation of this site was still detectable at low levels in *spo11-YF* strains indicating that it is not solely dependent on meiotic DNA break formation.

### Protein levels do not change in response to meiotic DNA breaks

We tested whether the meiotic DNA break response also affects protein levels. To this end, we quantified proteins in the same replicate cultures described above. This analysis yielded robust quantitative data for just under half of the yeast proteome (2,627 proteins). Measurements were highly correlated between replicates and samples (**Supplemental Figure 7A-C**). Accordingly, log_2_ fold changes in response to DNA break formation were narrowly centered on zero (**Supplemental Figure 8A**), indicating a surprisingly steady proteome during meiotic DNA break formation.

Only four proteins changed significantly in abundance between *SPO11* and *spo11-YF* cultures (adjusted p-value < 0.1): the levels of Rad51, Leu2, and Dbp2 were elevated in *SPO11* cells; Sml1 levels were decreased (**Figure 4A**). We validated the relative changes in Rad51, Sml1, and Dbp2 levels by immunoblotting (**Figure 4B-D**). Rad51 and Sml1 are known targets of the DNA damage response; Rad51 is a recombinase required for DNA break repair that is induced upon DNA damage and also protected from degradation (Basile et al., 1992; Woo et al., 2020), whereas the ribonuclease reductase RNR inhibitor Sml1 is degraded (Andreson et al., 2010; Zhao, 2001). The resulting changes in abundance are in agreement with our data (**Figure 4A-C**), indicating that these aspects of the DNA damage response remain active in response to meiotic DNA breaks. Differences in Leu2 levels were expected given that *SPO11* cells contained four copies of the *LEU2* gene whereas the *spo11-YF* cells only contained two (**Supplemental Table 1)**. The significantly higher Leu2 levels in the *SPO11* strain thus served as a good internal control. Dbp2 is an essential RNA-binding protein without known function during meiosis or the DNA damage response. When testing its meiosis-specific depletion, we observed only a minor loss in gamete viability (**Supplemental Figure 9A**). Taken together, the small overall change in protein levels indicates that the meiotic DNA break response does not involve substantial changes in translation or protein degradation. The robust proteomics data also indicated that the observed changes in protein phosphorylation were not driven by underlying changes in protein abundance (**Figure 4E**; **Supplemental Figure 8B**).

**Figure 4:**
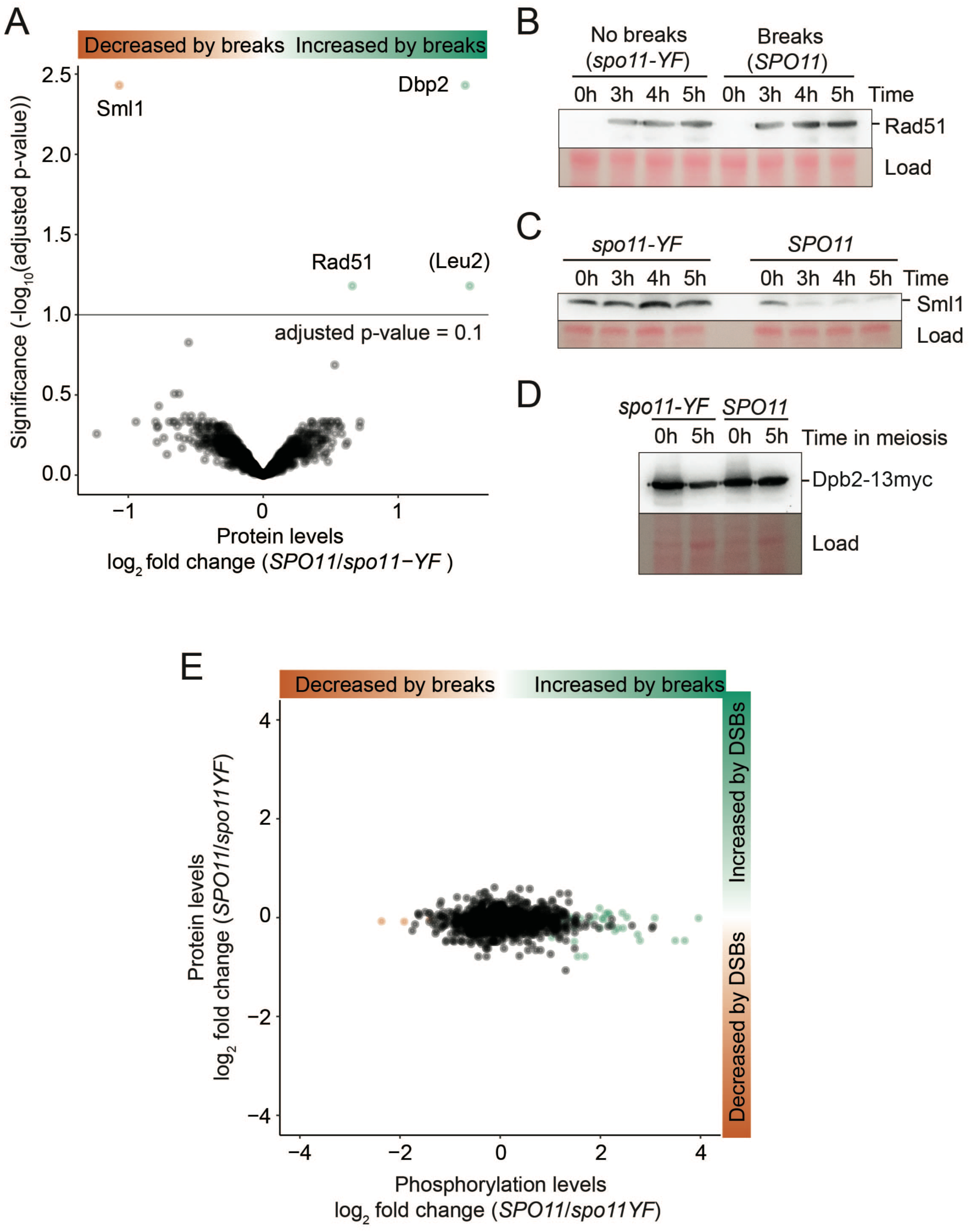
Only a few proteins change in abundance in response to meiotic DNA breaks. (A) A volcano plot showing differentially expressed proteins, each dot represents a protein. The y-axis shows -log_10_(Benjamini & Hochberg adjusted p-value) and the x-axis shows log_2_ fold change. (B) Western blot analysis of Rad51 in *spo11-YF* and *SPO11* cells during meiosis is shown. Ponceau S was used as the loading control. (C) Comparison of Sml1 levels in *spo11-*YF and *SPO11* cells during meiotic prophase. Sml1 was diminished after 3 hours into meiosis concomitantly with DNA break formation, Ponceau S staining of the membrane was the loading control. (D) Assessment of Dbp2 levels by western blotting prior to meiosis (0h) and during meiotic prophase (5h) is shown. Since there are no Dbp2 antibodies available, Dbp2 was tagged with a 13xmyc tag and anti-myc antibody was used for detection of Dbp2. Ponceau S was the the loading control. (E) Plot showing correlation between protein level log_2_ fold changes (y-axis) and phosphosite level log_2_ fold changes (x-axis). Each dot represents a phosphosite. While log_2_ fold changes for phosphosites distributed widely, protein log_2_ fold changes for proteins centered around 0.

### Activation of the signaling pathway for damage-dependent transcription

The robustness of protein abundances in response to meiotic DNA break formation was surprising, as it contrasted with the transcriptional changes that are characteristic of the canonical DNA damage response (Elledge & Davis, 1990; Gasch et al., 2001; Huang et al., 1998; Jaehnig et al., 2013; Tsaponina et al., 2011). We therefore investigated whether the effector kinases Rad53 and Dun1, which trigger the DNA-damage-dependent transcription changes, were indeed active during the response to meiotic DNA breaks. Western analysis of Rad53 in our experimental strains showed phosphorylated, slower migrating forms of Rad53 only in *SPO11* cells but not in *spo11-YF* cells (**Figure 5A**), indicating that Rad53 is activated by meiotic DNA breaks. In addition, we detected DNA break-specific phosphorylation of Dun1 at position S10 (Chen et al., 2007) (**Supplemental Table 3**) and reduction of Sml1 protein levels (**Figure 4A,C**), both of which are hallmarks of Dun1 activation as Sml1 is a Dun1 target (Zhao & Rothstein, 2002). Thus, key regulators of the transcriptional response to canonical DNA damage are activated also in response to meiotic DNA break formation.

**Figure 5:**
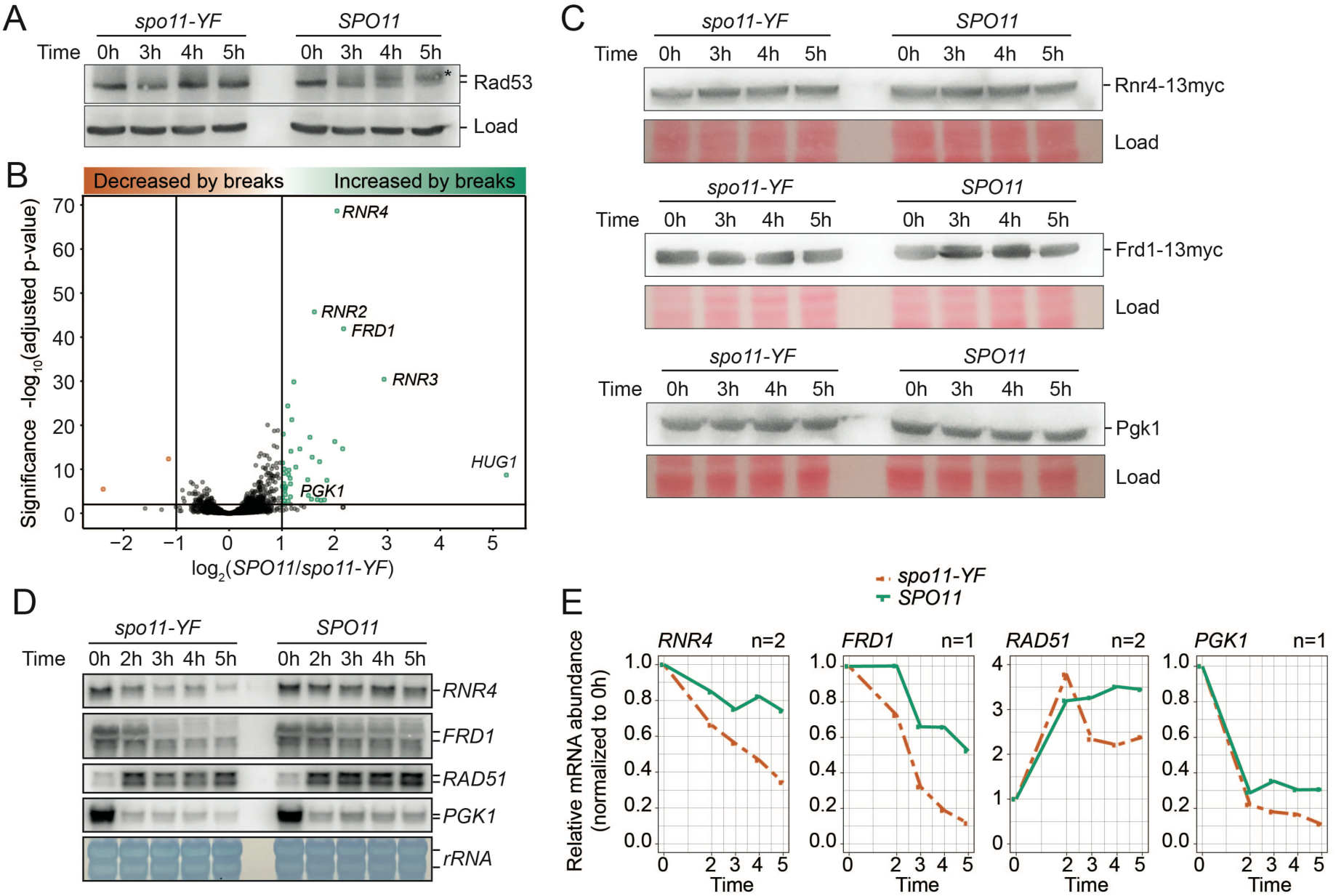
Activation of a transcriptional program in response to meiotic DNA breaks. (A) Western blot analysis of Rad53 in *spo11-YF* cells and *SPO11* cells. We used the mobility shift of Rad53 as a proxy for its phosphorylation and activation. A cross-reacting band produced by the Rad53 antibody was the loading control. Note: this analysis was performed on our experimental strains, which lack Pph3 phosphatase, resulting in more easily detectable Rad53 autophosphorylation than seen in strains with active Pph3 (Cartagena-Lirola et al., 2008; Falk et al., 2010). (B) A volcano plot showing the results of mRNA-Seq analyses of *spo11-YF* cells and *SPO11* cells. The y-axis shows -log_10_(Benjamini & Hochberg adjusted p-value) and the x-axis shows log_2_ fold change between *SPO11* and *spo11-YF*. Each dot represents a gene. (C) Western blot analysis of select transcriptionally induced genes. We used a 13xmyc tag to tag Rnr4 and Frd1 as antibodies for these genes were unavailable. Ponceau S staining of the membranes was the loading control. (D) Northern blot analysis of select transcriptionally induced genes. *RNR4, FRD1 and PGK1* were induced at the RNA level > 2 fold in *SPO11* cells compared to *spo11-YF* cells. *RAD51* was induced ∼ 1.6 fold. Loading was normalized according to RNA concentrations measured after RNA extraction. (E) Quantification of the Northern blots in (D) and additional replicate experiments. The plot shows the change of mRNA abundance compared to the 0-hour time point for each sample group (*SPO11* or *spo11-YF*). For n=2 panels, we used the average of two biological replicates. Data in this figure and Supplemental Figure S9 were from 4 independent time courses involving 3 different sets of *SPO11* and *spo11-YF* strains.

### A transcriptional response to meiotic DNA breaks

We tested if activated Rad53 and Dun1 induce transcription of DNA damage-response genes during meiosis. To this end, we conducted mRNA-seq experiments using the same strains as described above, comparing mRNA levels of *SPO11* and *spo11-YF* cultures. We found that 7% of genes (373 of 5,386) were differentially expressed (adjusted p-value < 0.01), and 42 genes (<1%) were upregulated more than two-fold in response to DNA break formation. This group contained many genes of the canonical gene expression response to DNA damage, including *RNR2*, *RNR3,* and *RNR4* (Elledge & Davis, 1987, 1990; Huang & Elledge, 1997) (**Figure 5B**).

We conducted several tests to confirm that, indeed, the proteome is highly fixed despite DNA break-dependent transcriptome changes. We verified that the discordance could not be explained by low coverage of the proteome data. Of the 287 transcriptionally upregulated genes, 165 (57%) were quantified in the proteome data, including 23 (54%) of the 42 genes with >2-fold change suggesting that the proteomics experiment captured a representative fraction of the proteome. None of these genes changed at the proteome level. Furthermore, immunoblotting of several DNA break-induced factors, i.e. Rnr4, Frd1, and Pgk1, throughout meiotic prophase confirmed that the protein levels were unaffected by meiotic DNA break formation and stable throughout meiosis both in *spo11-YF* and *SPO11* cells (**Figure 5C**) despite the increased levels of *RNR4*, *FRD1* and *PGK1* transcripts detected by mRNA-seq (**Figure 5B**).

### Meiotic entry is associated with strong reduction in mRNA abundance

To further investigate these unexpected results, we analyzed the transcript levels of several genes in a meiotic time course by Northern blotting. We observed two competing effects on mRNA levels. First, all analyzed transcripts experienced a noticeable drop in abundance as cells progressed through meiotic prophase (**Figure 5D, E**). For vegetatively expressed transcripts, like *RNR4*, *FRD1* and *PGK1*, overall mRNA abundance during meiosis dropped upon meiotic entry and remained at lower levels throughout the remainder of the time course (**Figure 5D,E**). This downregulation of *RNR4*, *FRD1, and PGK1* transcripts has also been observed in wild-type cells undergoing meiosis (**Supplemental Figure 9D**)(Cheng et al., 2018). Once cells progressed further into meiotic prophase, transcripts that were initially induced upon meiotic entry, such as *RAD51* and *HOP1* (**Figure 5D,E, Supplemental Figure 9B,C**), also dropped in abundance. The drops were observed regardless of whether samples were normalized by total cell number or total RNA (**Figure 5D,E, Supplemental Figure 9B,C**) and suggest that despite stable protein levels, transcripts levels drop when the cells enter meiotic prophase.

Second, as suggested by mRNA-seq results, the analyzed transcripts levels were higher in *SPO11* cells compared *spo11-YF* cells, confirming the induction of a transcriptional program in response to meiotic DNA breaks. However, the overall decrease in transcript abundance during meiosis overshadowed this elevation (**Figure 5D,E, Supplemental Figure 9B,C**). Taken together, mRNA and protein data suggest that protein levels for these genes are determined at or soon after meiotic entry when mRNA levels are highest, and later upregulation of these genes is not substantial enough to cause protein level changes.

## Discussion

In this study, we combined three systems-level approaches, i.e. transcriptomics, proteomics and phosphoproteomics, to capture the breadth of the meiotic DNA break response in *S. cerevisiae*. Our analyses identified 332 DNA break-dependent phosphorylation events, substantially expanding our current knowledge of the meiotic DNA break response. The breadth of detection also highlights the power of using complementary data acquisition techniques (DDA, DIA) and different analyses of the mass spectra (fold enrichment, presence/absence) for obtaining a large and high-quality dataset. Notably, the two approaches, fold enrichment and presence/absence, yielded qualitatively distinct groups of phosphorylation events. Fold enrichment analysis recovered many phosphorylation events that are not specific to meiosis, such as Hta1/2 S129 (γ-H2A) and Cbf1 S45, but are induced by different forms of canonical DNA-damage signaling (Cobb et al., 2005; Smolka et al., 2007). In comparison, presence/absence analysis recovered most of the known meiosis-specific phosphorylation events, including Zip1 S75, Hed1 T40 and Rad54 T132, which therefore appear to be regulated in an on/off-switch like manner in response to meiotic DNA breakage. Thus, the meiotic DNA break response elaborates on features of the canonical DNA damage response by adding a large number of targets that specifically respond to meiotic DNA breaks.

Our data set complements and expands on a published phosphoproteomics analysis that compared strains with active or inactive Mek1 kinase (Suhandynata et al., 2016) by identifying targets of all DNA break-dependent kinases and thus providing a comprehensive view of the phosphorylation events in response to DNA breaks. Importantly, our experimental setup also blocked cells from exiting prophase. As inactivation of Mek1 or Spo11 allows cells to prematurely exit from meiotic prophase (Suhandynata et al., 2016), phosphorylation differences between prophase and metaphase/anaphase can identify a substantial number of cell-cycle-dependent hits that are difficult to distinguish from the immediate meiotic DNA break response. Our analyses therefore provide a snapshot of the immediate DNA break response prior to exit from meiotic prophase.

Surprisingly, DNA break formation did not greatly impact the proteome of meiotic prophase. Our results suggest that this proteomic robustness arises from a combination of high protein stability and diminishing mRNA abundances, likely due to increased mRNA degradation. We show that many vegetatively expressed genes maintain their pre-meiotic protein levels throughout meiotic prophase, while their mRNA levels plummet after cells enter meiosis. Thus, although meiotic DNA breaks induce a clear transcriptional response, the increases in transcript levels were not large enough to alter the protein levels in a detectable manner.

The stability of protein abundances during meiotic prophase is consistent with the finding that most proteins are long-lived and that concentrations drop primarily as a consequence of dilution due to cell division (Martin-Perez & Villén, 2017), which does not occur during meiotic prophase. Targeted proteolysis via the meiosis-specific APC/Ama1 ubiquitin ligase or other mechanisms are important for meiotic prophase progression in yeast and mice (Kwon et al., 2003; Okaz et al., 2012), but the unchanged proteome observed here suggests that these mechanisms, while clearly active, do not become differently active in response meiotic DNA break formation. Rather, DNA break-dependent protein abundance changes were limited to a few select proteins with specialized modes of regulation, including Sml1, whose proteolysis is triggered by phosphorylation and subsequent ubiquitination (Zhao & Rothstein, 2002) and Rad51, which is protected from degradation by phosphorylation (Woo et al., 2020).

The drop in transcript levels as cells enter the meiotic program may be related to the unique aspects of mRNA metabolism in meiotic prophase. For example, N6-adenosine methylation of transcripts regulates both meiotic entry and meiotic commitment (Agarwala et al., 2012; Bushkin et al., 2019; Shah & Clancy, 1992). In addition, RNA stability is uncoupled from polyA-tail length during meiosis, likely through differential regulation of the RNA degradation protein Xrn1 (Wiener et al., 2021). Intriguingly, we observed DNA break dependent phosphorylation of Xrn1 on S1510. Although the effect of this phosphorylation on Xrn1 activity is unknown, it is possible that this phosphorylation event might reduce Xrn1 activity. A subsequent general slowdown in mRNA degradation during meiotic DNA breakage could explain why we observed not only the predicted induction of canonical DNA-damage response genes but also increased mRNA abundances of house-keeping genes, such as *PGK1*. Such potential interplay of the meiotic DNA break response with the mRNA degradation machinery warrants further analysis.

Taken together, our results suggest extensive rewiring of the canonical DNA-damage response in the context of meiotic DNA breaks by deemphasizing the role of proteome changes and instead exploiting and expanding post-translational signaling. We speculate that this shift toward posttranslational signaling reflects the unique needs created by the programmed induction of nearly 200 DNA breaks. As protein modifications have the crucial potential to create spatially distinct and constrained signals, they are uniquely suitable to support the patterning of the meiotic recombination landscape and the creation of local or chromosome-wide dependencies in a shared nuclear environment, which is a key feature of meiotic recombination (Kar & Hochwagen, 2021).

## Methods

### Yeast strains and meiotic time courses

All strains used in this study were derived from the SK1 background. **Supplemental Table 1** lists the genotypes of these strains. Meiotic time courses were set up by growing cells at room temperature (25°C) in rich medium (YPD) for ∼24 hours, followed by inoculation at a final OD_600_ of 0.3 in premeiotic BYTA medium (1% yeast extract, 2% bactotryptone, 1% potassium acetate, 50 mM potassium phtalate) and growth for 16-17 hours at 30°C. Cells were washed twice in sterile water and diluted to 1.9 OD_600_ in SPO medium (0.3% potassium acetate) and sporulated at 30°C. The time of resuspension in SPO was defined as the 0h time point.

### Flow cytometry analysis

We collected 150 μl of meiotic culture at the indicated time points and fixed cells with 350 μl 100% ethanol. The samples were stored at 4°C until further analysis. To prepare cells for flow cytometry, cell pellets were resuspended in 500 μl 50mM Na-Citrate and treated with 0.7 μl RNAse A (20-40 mg/ml stock; Sigma) at 50°C for at least 1 day. 5 μl Proteinase K (VWR, 20 mg/ml) was added to the samples and incubated at 50°C for at least 1 day before addition of 500 μl of 50 mM Na-Citrate with 0.1 μl SYTOX green (Invitrogen, 5 mM solution in DMSO). Prior to cytometry, samples were sonicated for approximately 5 seconds at 10% amplitude. Signal was collected using a BD Accuri C6 Flow cytometer.

### Preparation of protein samples for mass spectrometry

We collected 50 ml meiotic culture at 5 hours into meiosis, harvested cells at 4000 rpm for 3 minutes, and washed once with ice-cold sterile water at 4°C. Cell pellets were stored at -20°C until further processing. We used MS grade water in all the following steps and solutions. Proteins were extracted using an 8M urea lysis buffer (8M urea, 50 mM Tris-HCl, 75 mM NaCl, 1x Calbiochem protease inhibitor, 1 mM PMSF, 1x Thermo Fisher halt phosphatase and protease inhibitor). 2x cell pellet volume of lysis buffer and 1x cell pellet volume of acid washed glass beads (Millipore Sigma) were added to 1.5 ml tubes, and samples were agitated with Digital Vortex Mixer (Fisher Scientific) at 4°C 3 times for 10 minutes separated by 2-minute cooling intervals. The samples were centrifuged at 13,000 rpm for 20 minutes and the supernatants were transferred to new 1.5 ml tubes. Protein concentrations were measured using a Quick Start Bradford Protein Assay (Bio-Rad) and 200 μg of protein was processed for preparation of WCE protein samples.

Proteins were reduced in 5 mM dithiothreitol (DTT) at 37°C for 30 minutes and alkylated in 15mM iodoacetamide at room temperature (25° C) for 30 minutes in the dark. The alkylation was stopped by increasing the DTT concentration to 10 mM and incubating at room temperature for 15 minutes. Urea concentration was lowered by increasing the sample volume to 200 μl (7-fold dilution) with 50 mM Tris-HCl pH 8 solution. We added 3 μg Trypsin Gold (Promega) to each sample for digestion and incubated samples at 37°C overnight (approximately 16 hours) while shaking. Digestion was stopped by adding formic acid to a final concentration of 1%. Samples were dried under vacuum until all liquid was removed and resuspended in Buffer C (95% water, 5% acetonitrile, 0.1% formic acid). HyperSep tips (ThermoFisher Scientific) were used for clean-up following the kits instructions. Elution peptides were dried under vacuum until completely dry, resuspended in 100 µl amount of Buffer C and stored at -80° C. Peptide concentrations were determined using Pierce Quantitative Fluorometric Peptide Assay (ThermoFisher Scientific).

### Preparation of phospho-peptide enriched samples

We reduced 2,000 μg of protein samples and alkylated as described above. For phospho-peptide samples the volume was increased to 2,000 μl (7-fold dilution) and 30 μg Trypsin Gold (Promega) was added to each sample. After the samples were cleaned-up, phosphorylated peptides were enriched using the High-Select TiO_2_ Phosphopeptide Enrichment Kit (ThermoFisher Scientific) according to the manufacturer’s instructions.

### Mass spectrometry analysis for DDA data

Samples were analyzed using an EASY-nLC 1000 (ThermoFisher Scientific) coupled to a QEHF instrument (ThermoFisher Scientific). Peptides were separated using a PepMap C18 column (Thermo Fisher Scientific) with 155 minute gradient of Buffer A (0.1% Formic Acid) and Buffer B (80% Acetonitrile, 0.1% Formic Acid). Full MS spectra were collected in a scan range of 375-1500 with resolution of 120,000. AGC target was set to 3e6 and top 20 peptides were selected for further analysis with an isolation window of 1.5 m/z with a maximum injection time of 100 m/s. MS2 spectra were collected with a resolution of 30,000 and AGC target 2e5 with an isolation window of 1.5 m/z and normalized collision energy (NCE) of 27 in centroid mode.

### Mass spectrometry analysis for DIA data

For DIA runs, a full MS scan was collected with a resolution of 120,000 and AGC target of 3e6 between 350-1650 m/z. Each full MS scan was followed by 24 DIA windows with a resolution of 60,000 and AGC target of 1e6. Maximum injection time was set to auto, and normalized collision energy (NCE) was set to 27 in profile mode. The DIA windows are provided in **Supplemental Table 2**.

### Data analysis of DDA data

Raw data were processed in MaxQuant (version 1.5.5.1)(Tyanova et al., 2016) using the proteome of *Saccharomyces cerevisiae* strain S288C (downloaded from Uniprot on August 8 2017) with default settings. For phospho-enriched samples Phospho(STY) was selected as a variable modification. Data was further analyzed and graphed in custom made R scripts using packages tidyverse, ggplot2 and limma (Ritchie et al., 2015; Wickham, 2016; Wickham et al., 2019). Proteins from contamination and reverse search were filtered out. For protein level data, LFQ intensities were first normalized to parts per million and log_2_ transformed. Log_2_ transformed values were tested for significance analysis using limma. For analysis of phospho-proteome changes, we used the Phospho(STY) file. Phosphorylation level data was filtered out to only include phosphosites with 0.9 localization probability. Then the phosphorylation site intensities were normalized to parts per million and log_2_ transformed before significance analysis with limma.

### Data Analysis of DIA data

DIA data was analyzed using Spectronaut (v 13.8.190930.43655) against a project-specific spectral library (phospho-DDA data). Perseus plug-in Peptide Collapse program was used to convert the Spectronaut (Biognosys) report file to intensity values at the peptide level (Bekker-Jensen et al., 2020). Peptide level intensity values were normalized to parts per million and log_2_ transformed before performing significance analysis using limma.

### GSEA and motif analysis

Gene set enrichment analysis was performed using R package clusterProfiler using the enrichGO function (Yu et al., 2012). Motif analysis was performed with sequences of 3 amino acids on N and C terminal sides around the phospho-sites using rmotif-x with p-value cut-off of0.05 (Wagih et al., 2016). Sequences of all phosphosites detected in our study were used as the background dataset.

### Preparation of mRNA-Seq samples

#### RNA Extraction

We harvested 1.6 ml of meiotic culture at 4 hours into meiosis and centrifuged samples at 3000 rpm for 5 minutes at 4°C. All supernatant was removed and the pellet was resuspended with 1 mL of Tris-EDTA (10 mM Tris pH 8, 1 mM EDTA) buffer. The samples were spun down again and the supernatant was removed. The samples were stored at -80° C until further processing. RNA extraction was performed using the Rneasy Mini Kit (Qiagen). 600 µl of RLT buffer with 1% (v/v) β-mercaptoethanol and ∼200 mg glass beads were added to the pellets. The samples were agitated for 20 minutes at 4°C and spun down at 14,000 rpm for 2 minutes. Supernatant was transferred to a new microcentrifuge tube and mixed 1:1 with 70% ethanol. Samples were transferred to Rneasy columns and RNA extraction was completed following the kit’s instructions. We measured the RNA integrity using Agilent RNA screentape and RNA concentration using Qubit RNA HS assay kit (ThermoFisher Scientific). We used 1.2 µg total RNA for mRNA purification. mRNAs were purified using Sera-Mag oligo(dt) magnetic particles (Sigma-Aldrich). mRNAs were fragmented with Ambion mRNA fragmentation buffer and fragmented mRNAs were purified using RNeasy MinElute Kit (Qiagen). Final elution volume was 9µl.

#### First- and Second-Strand Synthesis

First-strand synthesis was performed in a similar manner as described in (Parkhomchuk et al., 2009). For first-strand synthesis, 8 µl fragmented mRNAs were mixed with 1 µl of random hexamers (Invitrogen) and 1 µl of 10 mM dNTPs. The samples were incubated at 65°C for 5 min and chilled on ice for 1 minute. We added 10 µl of a master mix with final concentrations of 1X RT Buffer (ThermoFisher Scientific), 10 mM MgCl_2_, 20 mM DTT, 4 U/µl RnaseOUT (ThermoFisher Scientific) and 20 U/µl of SuperScript III RT (ThermoFisher Scientific) to the RNA samples. The samples were then first incubated at 25°C for 10 minutes, followed by a 50-minute incubation at 50°C. The reaction was stopped by incubating samples at 75°C for 15 minutes. For dNTP cleanup, 80 µl water, 1 µl glycogen, 10 µl 3 M NaOAc (pH 5.2), and 200 µl cold ethanol were added to the samples. Samples were stored at -80°C for 3-7 days. Samples were centrifuged at 14,000 rpm for 20 minutes at 4°C. Supernatant was removed and 500 µl of cold 75% ethanol was added to the samples. Samples were centrifuged again at 14,000 rpm for 10 minutes at 4°C. Supernatant was removed and samples were resuspended in a mixture composed of 51 µl RNAse-free water, 1 µl of 10X RT buffer, 1 µl 100mM DTT, 2 µl of 25 mM MgCl_2_. Second-strand synthesis was performed as described in (Parkhomchuk et al., 2009).

#### Library Preparation

Library preparation was performed using TruSeq Library prep kit v1, but the adapters were used at 1:20 and 1U of UNG enzyme (Thermo Fisher Scientific) was added before PCR amplification to digest uridine containing templates to produce directional libraries. For uridine digestion, the samples were incubated at 37°C for 15 minutes and the reaction was terminated by incubation at 98°C for 10 minutes. Amplified DNA was run on a 1.5% agarose gel and DNA between 250 bp and 600 bp was extracted using a Qiagen Gel Extraction kit with a MiniElute column. DNA concentrations were measured using Qubit dsDNA HS Assay Kit (ThermoFisher Scientific) and KAPA library quantification kit (Roche). DNA sizes were checked with Agilent High Sensitivity D1000 ScreenTape. 75-bp pair-ended sequencing was performed on a NextSeq 500 instrument.

### Analysis of RNA-Seq samples

RNA-Seq reads were mapped to the SK1 genome using the nf-core RNA-Seq pipeline (Ewels et al., 2020; Patel et al., 2021; Yue et al., 2017). We used the salmon.merged.gene_counts.rds file from salmon output for further analysis. Combat-Seq was used for batch correction and DeSeq2 was used for principal components analysis and for differential gene expression analysis (Love et al., 2014; Zhang et al., 2020).

### Immunoblotting

For immunoblotting, 5 ml samples were collected at the indicated time points. The samples were spun down at 2,500 rpm for 2.5 minutes, and pellets were resuspended in 5% TCA. The samples were kept on ice for at least 10 minutes after resuspension. The samples were washed with 500 µl 1 M Tris, and resuspended in 80 µl of TE+DTT (0.8XTE, 200 mM DTT) buffer. After the addition of 30 µl 5x SDS buffer (190 mM Tris-acetate, 6% β-mercaptoethanol, 30% glycerol, 20% SDS, 0.05% bromophenol blue), the samples were incubated at 100°C for 5 minutes and stored at -80°C immediately. Proteins were run in handcast 10% 29:1 (acrylamide: bis-acrylamide) gels for Hrr25 blots and handcast 8% 29:1 (acrylimide: bis-acrylimide) gels for Rad53 blots. 4%-20% gradient gels (Bio-Rad) were used for Sml1 blots and 4%-15% gradient gels (Bio-Rad) were used for Frd1-13myc, Rnr4-13myc, and Rad51 blots. All blots were blocked with 5% milk. Primary antibodies were used at the following concentrations; Hrr25 ph-S438 (rabbit, 1:1000), Rad53 (goat, yc-19 Santa Cruz) at 1:500, Sml1(rabbit, AgriSera) at 1:1000, β-actin (rabbit, CST) at 1:1000, Myc-tag at 1:1000 (rabbit, CST). Anti-rabbit secondary antibody (Kindle Biosciences) was used at 1:2000, anti-goat secondary antibody (Kindle Biosciences) was used at 1:1000. The blots were visualized using KwikQuant Imager (Kindle Biosciences). The phospho-specific Hrr25 ph-S438 antibody was raised by Covance against the synthetic target peptide Ac-QQRD(pS)QEQQC-amide.

### Northern blotting

For Northern blotting, 6 ml samples were collected at the indicated time points, spun down at 2,500 rpm for 2.5 minutes and stored in 2 ml microcentrifuge tubes at -80°C immediately. Pellets were overlaid with 350 µl acid phenol chloroform pH4.5 (Thermo Fisher Scientific). After addition of 100 mg glass beads and 350 µl RNA buffer 1 (300 mM NaCl, 10 mM TrisHCl pH 6.8, 1mM EDTA, 0.2% SDS) samples were agitated at 4°C for 10 minutes in a Disruptor Genie (Scientific Instruments). Phases were separated by centrifuging 10 minutes at 14,000 rpm at 4°C and 300 µl of the aqueous phase were precipitated in 1 ml cold 100% ethanol at 4°C for 10 minutes. RNA was collected by centrifuging for 5 minutes at 14,000 rpm at 4°C and pellets were resuspended in RNA buffer 2 (10 mM TrisHCl pH 6.8, 1 mM EDTA, 0.2% SDS) at 65°C for 20 minutes before storing at at -20°C. RNA concentration was determined using a NanoDrop instrument. Samples were denatured for 10 minutes at 65°C in denaturation mix (40 mM MOPS pH 7.0, 50% formamide, 6.5% formaldehyde) and separated in a 1.1% agarose gel containing 6.2% formaldehyde and 40 mM MOPS pH 7.0. RNA was blotted onto a HybondN+ membrane using neutral transfer in 10x SSC and UV crosslinked. Radioactive probes were synthesized from gel-purified templates using a Prime-it RmT Random Labeling Kit (Agilent) and alpha-^32^P-dCTP (Perkin Elmer). Templates were produced by PCR using the following primers (*RNR4*: F 5’-cag ccg tag att cgt gat gtt ccc -3’, R 5’- gcg gac tta gac atg tca ctg gcc -3’; *FRD1*: F 5’- ggt ttg gcc ggg ctg gct gc -3’, R 5’- gca taa ttg ggc gac agt gat tgg -3’; *RAD51*: F 5’- cag ctt cag tac ggg aac ggt tcg -3’, R 5’- gcc ata cca cca tca act tgg gcg -3’; *HOP1*: F 5’- ccc aat ccc tgg aac ctt tac ccc -3’, R 5’- gct cct gta ggg ttg acg acg gag -3’; *PGK1*: F 5’- tga ctt caa cgt ccc att gga cgg -3’, R 5’- aac acc tgg acc gt cca gac -3’). Signals were measured using a Typhoon FLA9000 instrument.

### Data Availability

All mass spectrometry data are deposited at the PRIDE databases PXD031779 and PXD031781. RNA-seq data are deposited at the GEO database GSE197022.

## Acknowledgements

AH acknowledges funding by the US National Institutes of Health (R01GM11171 and R01GM123035). CV acknowledges funding by the US National Institutes of Health (R35GM127089 and 75N93019C00052/NH/NIH HHS/United States). We thank the NYU Department of Biology Sequencing Core for technical assistance and data processing.

## Supplementary Figures

**Figure S1:**
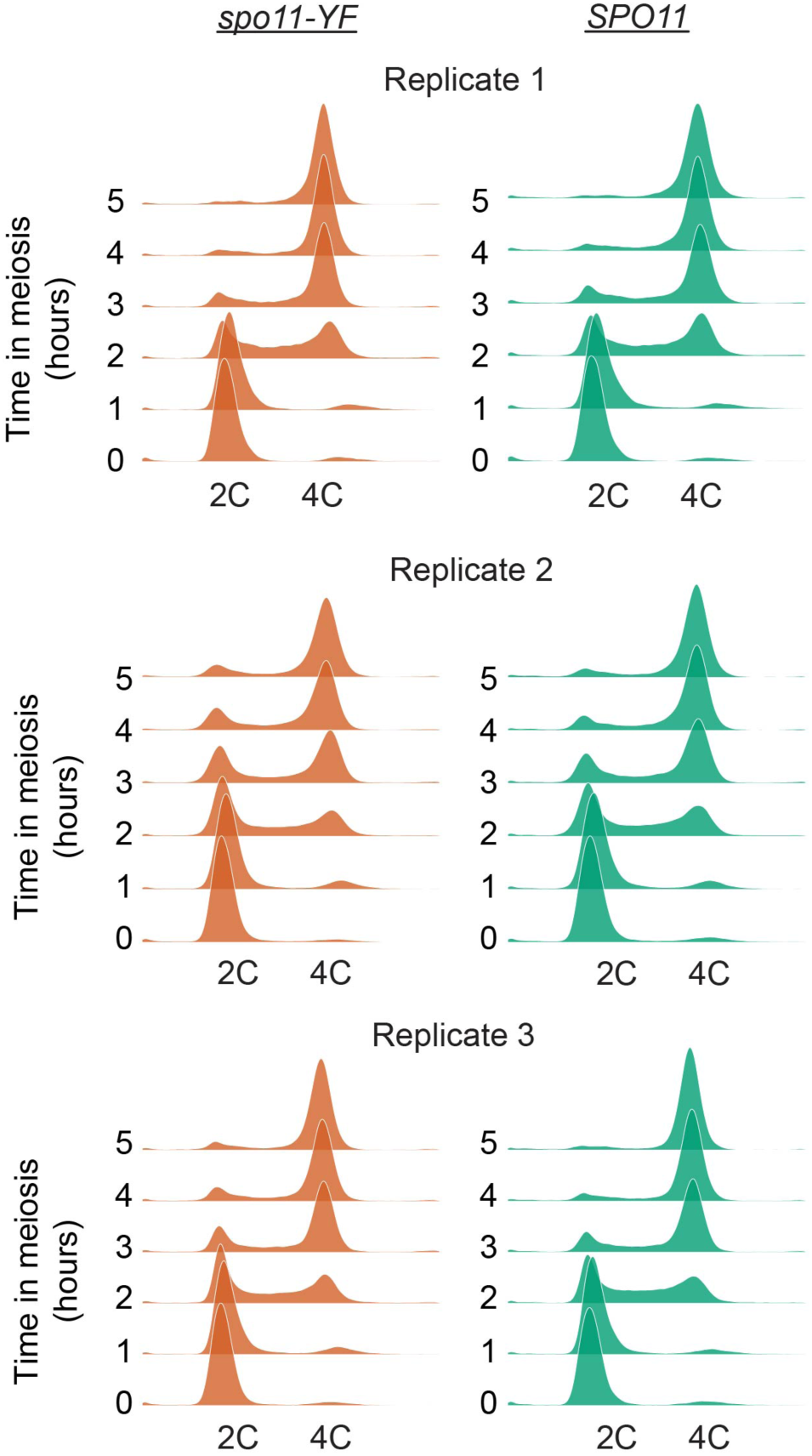
Synchrony of premeiotic DNA replication of biological replicates. We used DNA replication timing as a proxy for meiotic synchrony and meiotic progression. Flow cytometry results using SYTOX Green are shown for all three replicates.

**Figure S2:**
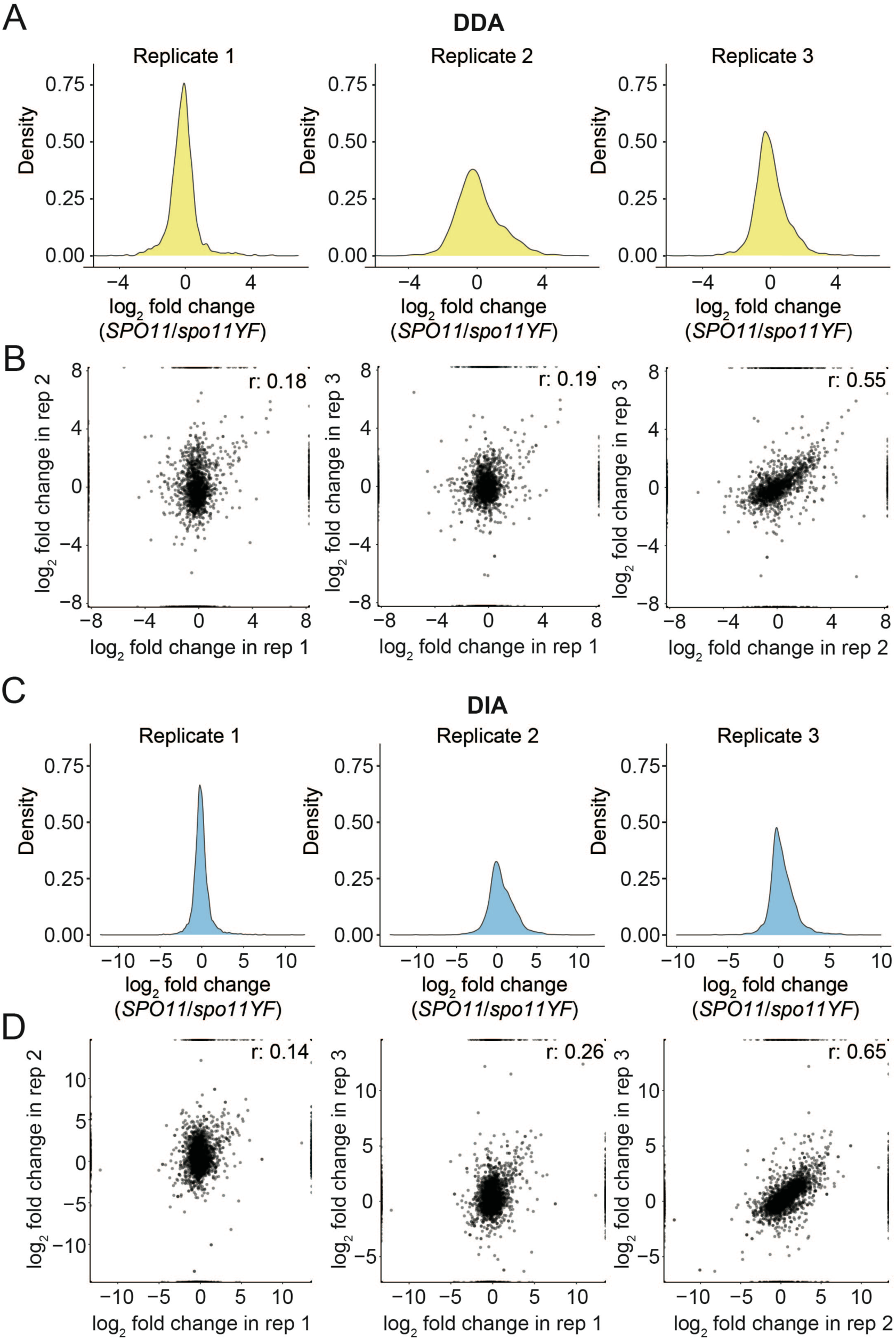
Assessment of DDA and DIA phosphoproteomics data. (A) Density plots for each replicate of the DDA phosphoproteomics dataset. log_2_(*SPO11 /spo11-YF*) values are on the x-axis and density of the values are on the y-axis. (B) Correlation of log_2_(*SPO11 /spo11-YF*) values between each replicate is shown for DDA data, each dot represents a phosphosite. (C) Density plots for each replicate of the DIA phosphoproteomics dataset. log_2_(*SPO11 /spo11-YF*) values are on the x-axis and density of the values are on the y-axis. (D) Correlation of log_2_(*SPO11 /spo11-YF*) values between each replicate is shown for DIA data, each dot represents a phosphopeptide.

**Figure S3:**
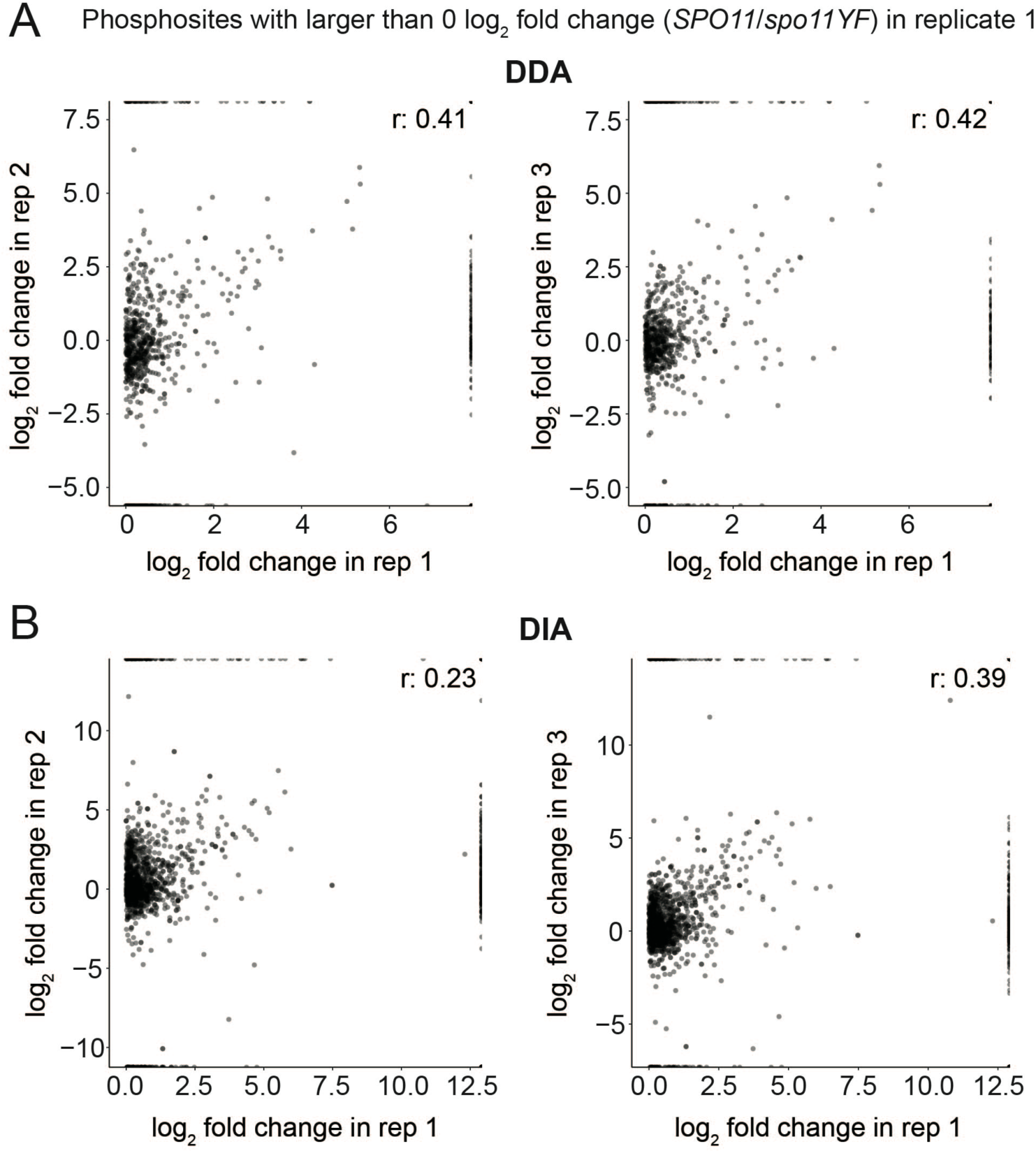
Further inspection of phosphoproteomics data from Replicate 1. (A) Correlation of log_2_(*SPO11 /spo11-YF*) values only for phosphosites with a log_2_(*SPO11 /spo11-YF*) value higher than 0 in replicate 1 is shown. These graphs are made using DDA data. Each dot represents a phosphosite. (B) Same as A, but for DIA data.

**Figure S4:**
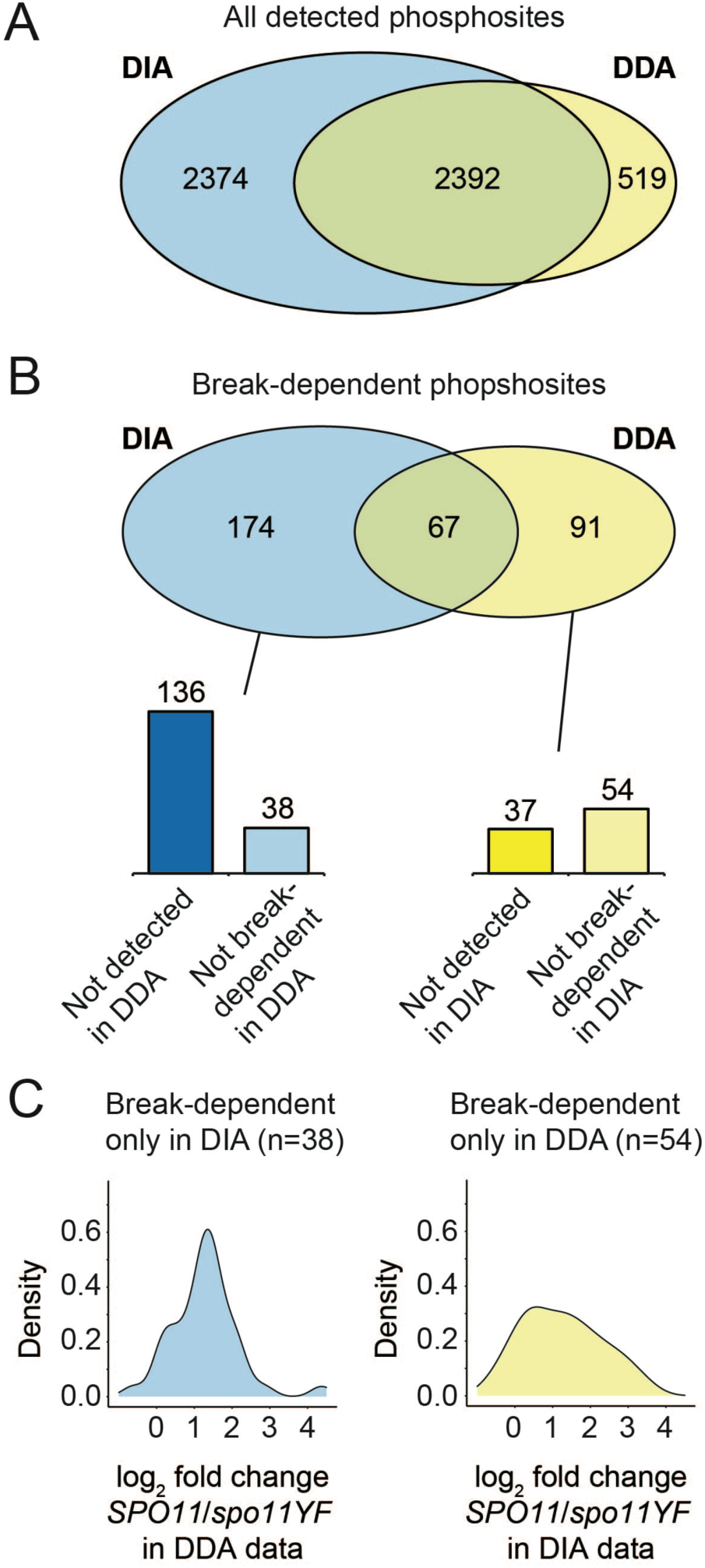
Differences and commonalities of DDA and DIA datasets. (A) A Venn diagram showing the overlap between DDA and DIA phosphosites for all phosphosites detected in at least 3 samples out of 6 in DDA or DIA datasets. (B) Same as A, but for DNA break-dependent phosphosites. While 67 sites overlap between DDA and DIA, there are in total 265 method-specific sites. Closer inspection revealed that 136 of 174 DIA-specific DNA break-dependent sites were not reliably detected in the DDA data (not present in at least 3 samples out of 6), indicating that these sites are missing from the DDA data due to missing values. The remaining 38 DIA-specific sites were reliably detected in DDA data, but were not categorized as DNA break-dependent after differential presence analysis. Similarly, 37 of 91 DDA-specific sites were not detected reliably in the DIA data. (C) The distribution of log_2_ fold changes (*SPO11*/*spo11-YF*) for 38 DIA-specific DNA break-dependent in the DDA data is between ∼ -0.6 and ∼4.3, and the distribution was centered around 1 (left panel).Only one site had a log_2_ fold change smaller than -0.5 (∼1.4 fold decrease) while 31 sites had log_2_ fold changes larger than 0.5 (∼1.4 fold increase), indicating that these sites trended largely correctly but were below the significance cutoff in the DDA data. The 54 DDA-specific DNA break-dependent sites had a distribution of log_2_ fold changes between ∼-0.7 and ∼3.5 in the DIA data, with the distribution being centered around 1 (right panel). While only one site had a log_2_ fold change smaller than -0.5 (∼1.4 fold decrease), 37 sites had log_2_ fold changes larger than 0.5 (∼1.4 fold increase). Since a trend of being induced in response to DNA breaks (log_2_ fold change > 0) is observed for the majority of the method-specific sites, we merged all the DDA and DIA datasets.

**Figure S5:**
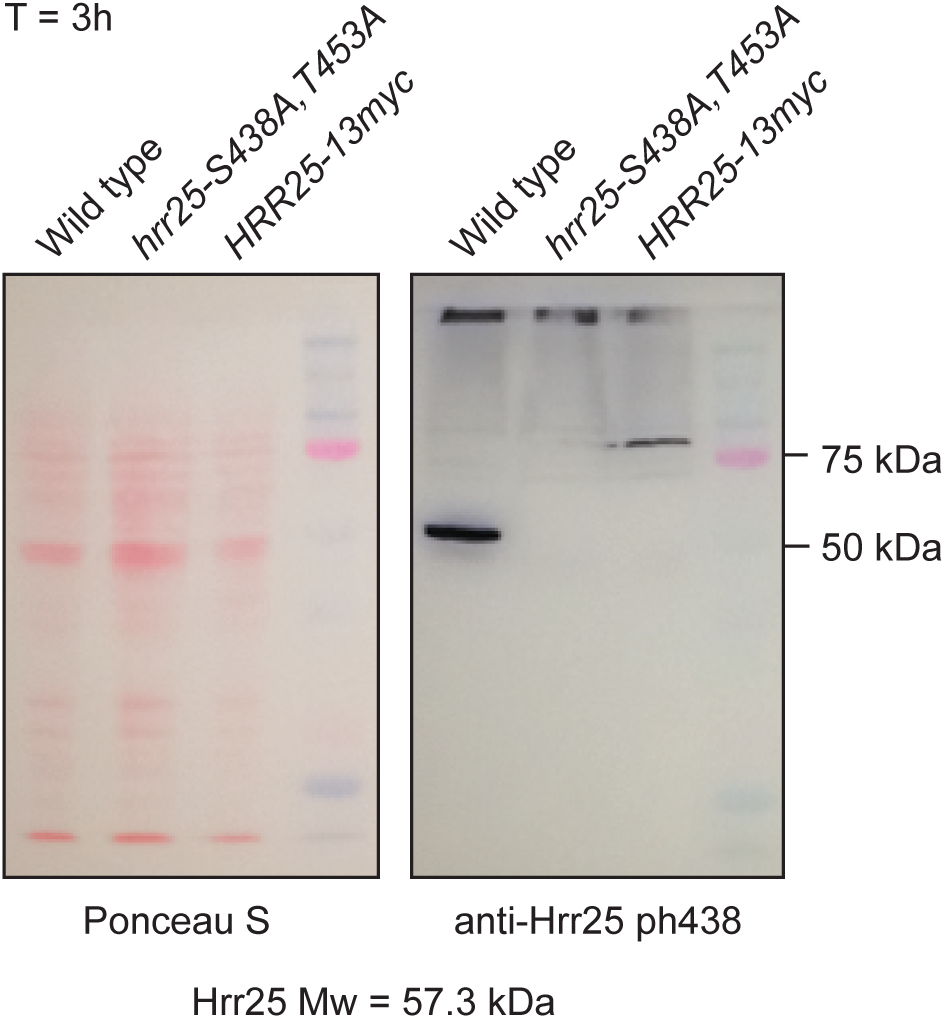
Specificity of Hrr25 phS438 antibody. (A) Western blot depicting the specificity of Hrr25 phS438 antibody. No signal was detected in non-phosphoryable S438A mutant, and a shift to a higher molecular weight was detected when Hrr25 was tagged with 13xmyc.

**Figure S6:**
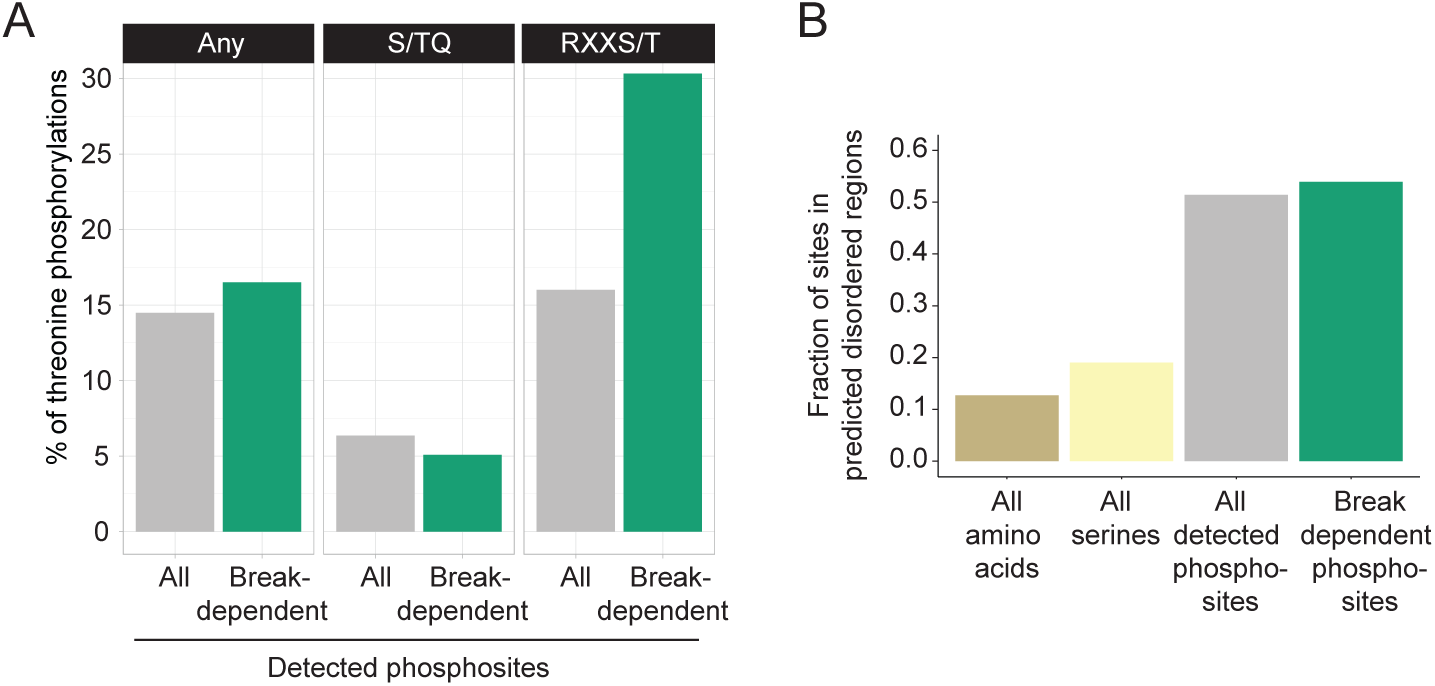
Other features of DNA break-dependent phosphosites. (A) Bar graph showing enrichment of threonines as the phosphorylated amino acid in DNA break-dependent Rxx<u>S/T</u> phosphorylation events, compared to all RxxS/T phosphorylation events detected in our data. No such effect is seen for DNA break-dependent <u>S/T</u>Q phosphorylation sites. (B) Bar graph showing the fraction of the detected phosphorylation sites that localize to IUPRED predicted disordered regions larger than 30 amino acids. Fraction of all serines in the yeast proteome that localize to these regions and fraction of the proteome in these regions are also shown for comparison.

**Figure S7:**
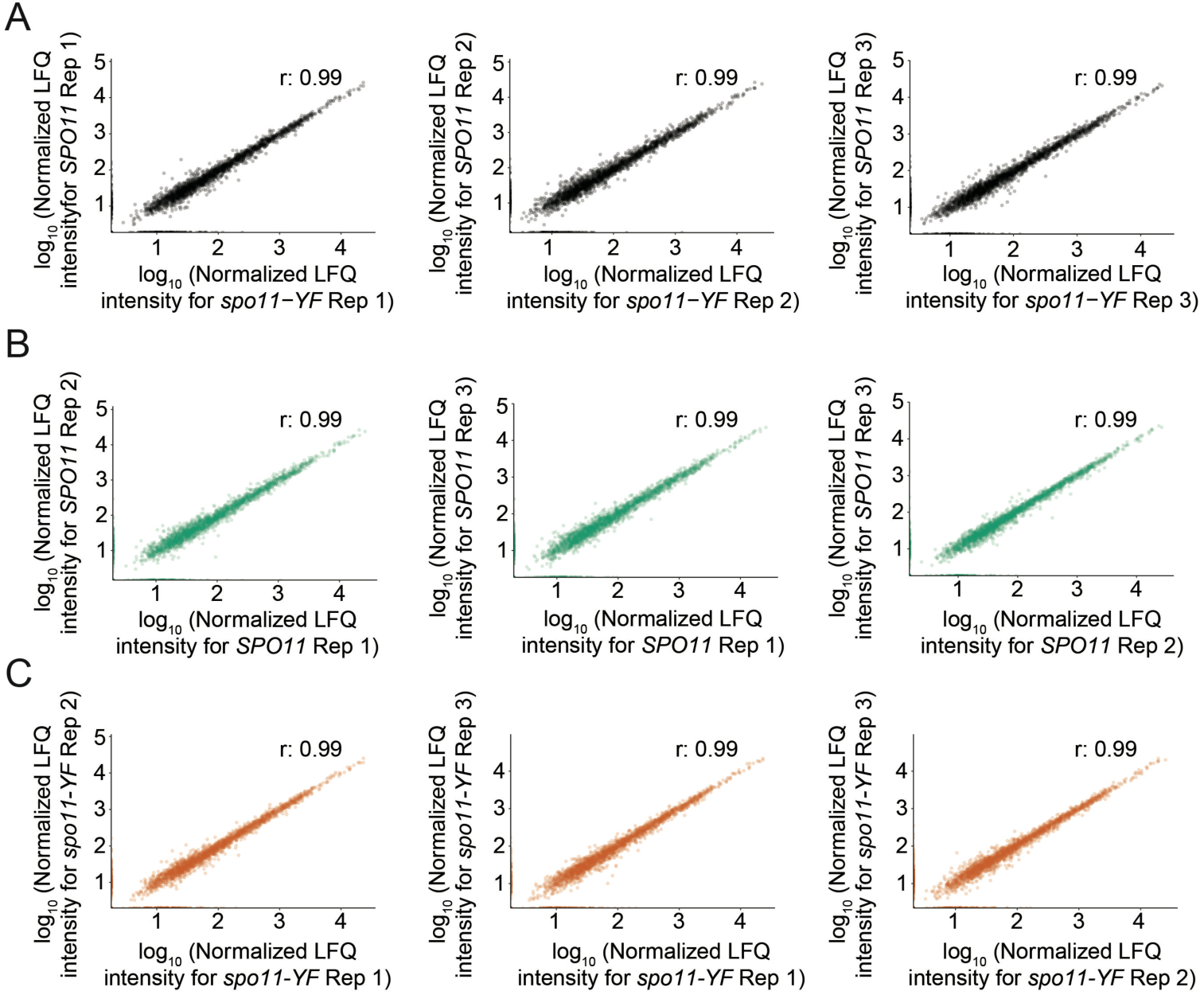
Assessment of proteomics data. (A) Scatter plots depicting the correlation between *SPO11* and *spo11-YF* samples within each replicate groups. (B) Scatter plots depicting the correlation of each *SPO11* sample with other *SPO11* replicate samples. (C) Scatter plots depicting the correlation of each *spo11-YF* sample with other *spo11-YF* replicate samples.

**Figure S8:**
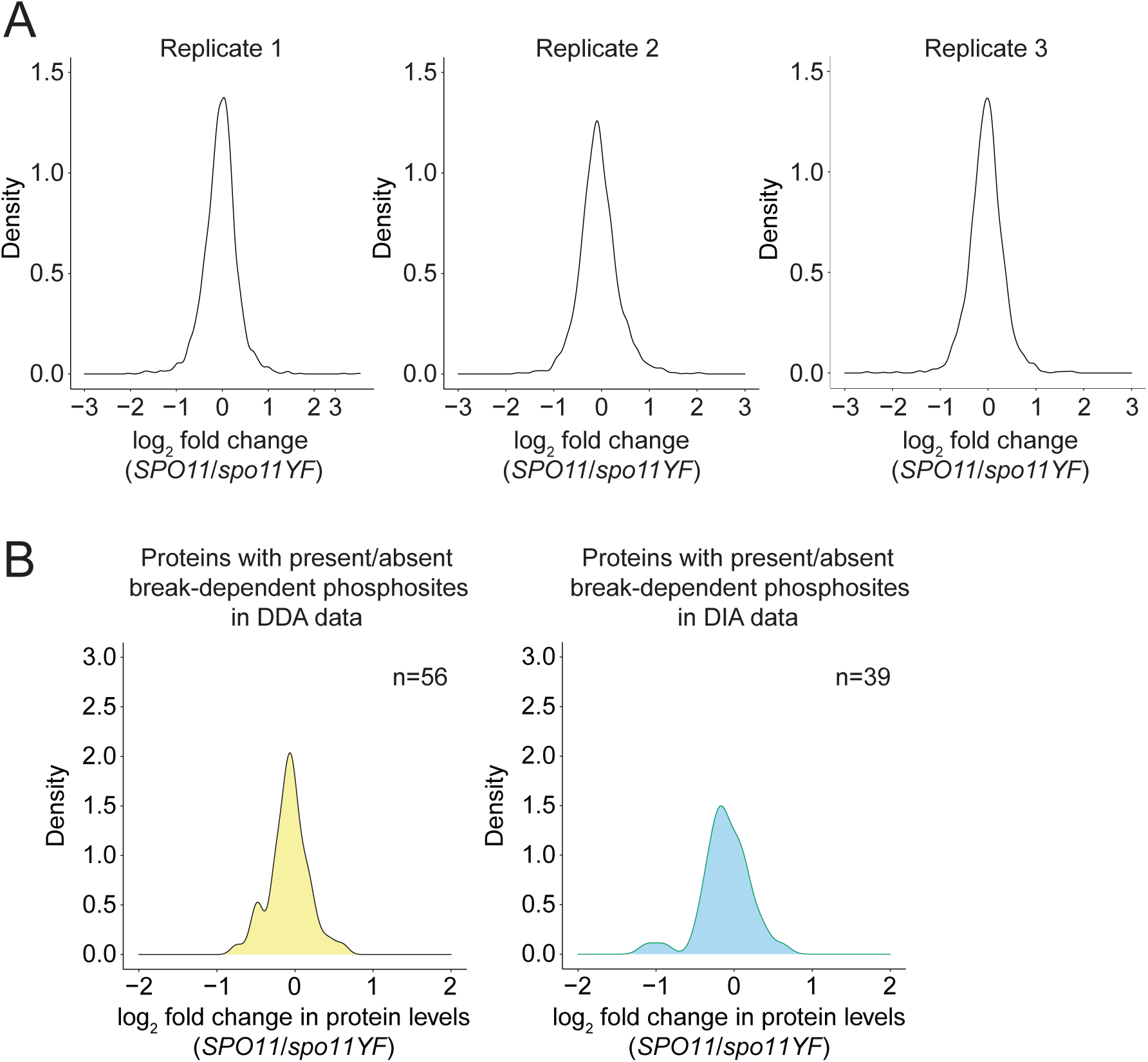
Distribution of protein level changes. (A) Density plots for each replicate in the proteomics dataset. log_2_(*SPO11/spo11-YF*) values are on the x-axis and density of the values are on the y-axis. (B) Density plots for proteins with qualitative (present/absent) DNA break-dependent sites in DDA data (left), and DIA data (right).

**Figure S9:**
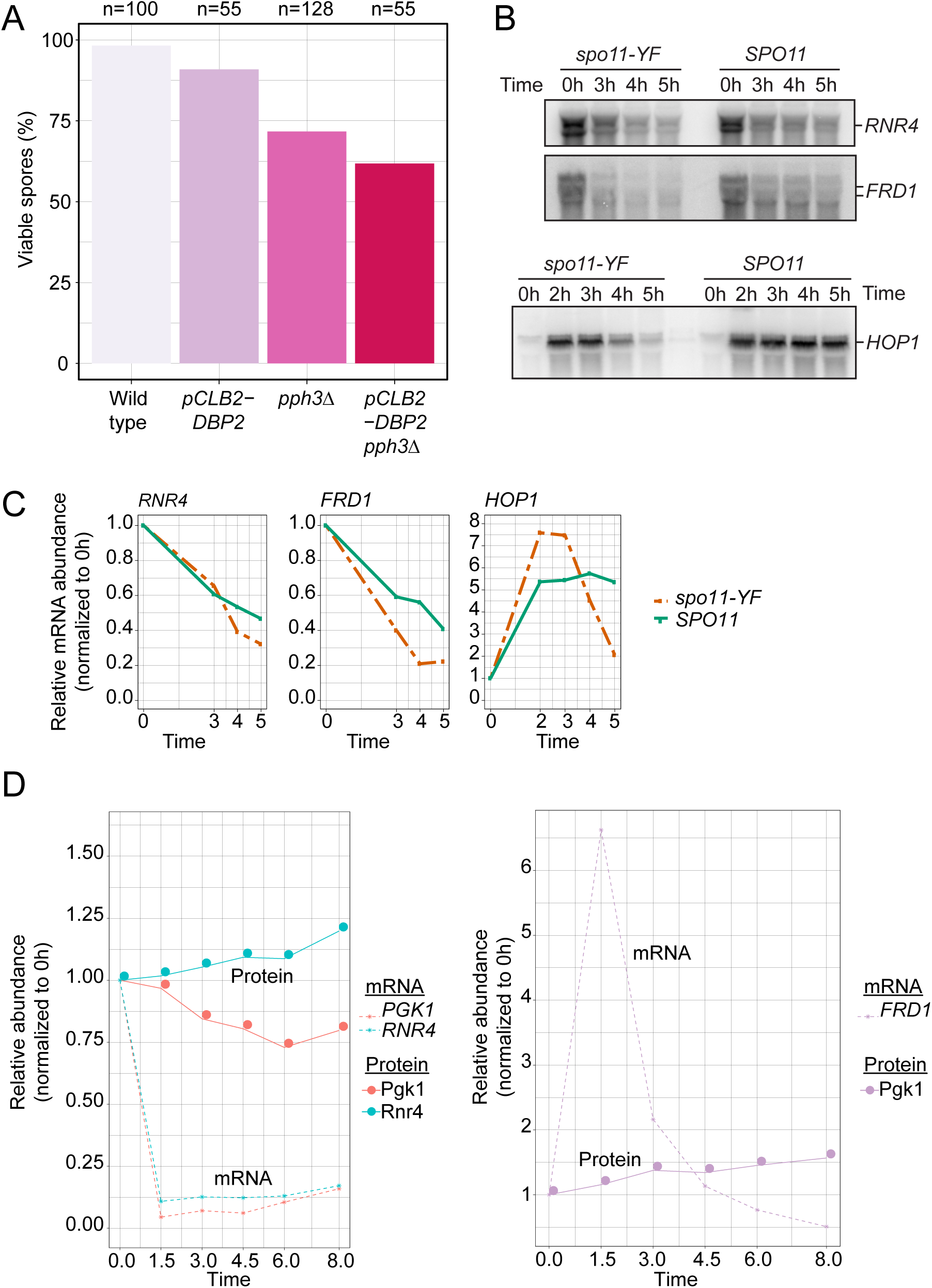
Effect of Dbp2 depletion on gamete viability, and dynamics of transcript and protein levels during meiosis. (A) Bar graph showing spore viabilities for wild type, *pph3Δ*, *pCLB2-DBP2*, and *pph3Δ pCLB2-DBP2* strains. (B) Northern blots for *RNR4, FRD1 and HOP1* in *SPO11* and *spo11-YF* cells. Loading is normalized based on number of cells (same culture volume is collected for all samples, OD = 1.9). (C) Quantification of the Northern blots in (B). Intensity values are normalized by dividing them to value at 0h. (D) Left panel; change in *RNR4* and *PGK1* transcript and protein levels throughout meiosis compared to 0h in wild-type cells, same for *FRD1* on the right. *FRD1* is plotted separately from *PGK1* and *RNR4* due to the different scale of its y-axis. Data is from the **Supplemental table 3** from (Cheng et al., 2018). We used the rpkm values from the indicated time points for plotting mRNA dynamics. For protein level changes, we used average of the two replicates from “total norm” columns.

---

**Supplemental Table 1:** Genotypes of the strains used in this study

| Strain | Genotype | Background | Figure |
| --- | --- | --- | --- |
| H10355 | MATa, ho::LYS2, lys2, HIS, URA, spo11-Y135F-HA-URA3, pph3Δ::LEU2, ndt80Δ::TRP1<br>MATalpha, ho::LYS2, lys2, HIS, URA, TRP1, spo11-Y135F-HA-URA3, pph3Δ::LEU2, ndt80Δ::TRP1 | SK1 | 2a,2b,<br>2d,4a,4b,4e,5a,5b,5c,5d,5d,<br>S1,S2,S3,S7,S8,S9B,S9C |
| H10423 | MATalpha, ho::LYS2, lys2, HIS, URA, LEU2, pph3Δ::LEU2, ndt80Δ::TRP1<br>MATa, ho::LYS2, lys2, HIS, URA, LEU, ndt80Δ::TRP1, pph3Δ::LEU2 | SK1 | 2a,2b,<br>2d,4a,4b,4e,5a,5b,5c,5d,5d,<br>S1,S2,S3,S7,S8,S9B,S9C |
| H11604 | MATa, ho::LYS2, lys2, ura3, leu2::hisG, HIS, trp1::hisG, spo11-Y135F-HA-URA3, pph3Δ::LEU2, ndt80Δ::TRP1, FRD1-13myc::HIS3MX6<br>MATalpha, ho::LYS2, lys2, ura3, leu2::hisG, HIS?, trp1::hisG, spo11-Y135F-HA-URA3, pph3Δ::LEU2, ndt80Δ::TRP1, FRD1-13myc::HIS3MX6 | SK1 | 5c, 5d, 5e, S9B, S9C |
| H11605 | MATa, ho::LYS2, lys2, ura3, leu2::hisG, HIS, trp1::hisG, spo11-Y135F-HA-URA3, pph3Δ::LEU2, ndt80Δ::TRP1, FRD1-13myc::HIS3MX6<br>MATalpha, ho::LYS2, lys2, ura3, leu2::hisG, HIS, trp1::hisG, pph3Δ::LEU2, ndt80Δ::TRP1, FRD1-13myc::HIS3MX6. | SK1 | 5c, 5d, 5e, S9B, S9C |
| H11641 | MATa, ho::LYS2, lys2, ura3, leu2::hisG, his3::hisG, trp1::hisG, DBP2-13myc::HIS3MX6, pph3Δ::LEU2, ndt80Δ::TRP1, spo11-Y135F-HA-URA3<br>MATalpha, ho::LYS2, lys2, ura3, leu2::hisG, his3::hisG, trp1::hisG, DBP2-13myc::HIS3MX6, pph3Δ::LEU2, ndt80Δ::TRP1 | SK1 | 4d |
| H11642 | MATa, ho::LYS2, lys2, ura3, leu2::hisG, his3::hisG, trp1::hisG, DBP2-13myc::HIS3MX6, pph3Δ::LEU2, ndt80Δ::TRP1, spo11-Y135F-HA-URA3<br>MATalpha, ho::LYS2, lys2, ura3, leu2::hisG, his3::hisG, his4X,<br>trp1::hisG,<br>DBP2-13myc::HIS3MX6,<br>pph3Δ::LEU2, ndt80Δ::TRP1, spo11-Y135F-HA-URA3 | SK1 | 4d |
| H11746 | MATalpha, ho::LYS2, lys2, ura3,<br>leu2::hisG, his3::hisG, trp1::hisG,<br>pph3Δ::LEU2, spo11-Y135F-HA-URA3, ndt80Δ::TRP1, RNR4-13myc::HIS3MX6<br>MATa, ho::LYS2, lys2, ura3,<br>leu2::hisG, his3::hisG, trp1::hisG,<br>pph3Δ::LEU2, spo11-Y135F-HA-URA3, ndt80Δ::TRP1, RNR4-13myc::HIS3MX6 | SK1 | 5c, 5d, 5e, S9B, S9C |
| H11747 | MATalpha, ho::LYS2, lys2, ura3,<br>leu2::hisG, his3::hisG, trp1::hisG,<br>pph3Δ::LEU2, ndt80Δ::TRP1, RNR4-13myc::HIS3MX6<br>MATa, ho::LYS2, lys2, ura3,<br>leu2::hisG, his3::hisG, trp1::hisG,<br>pph3Δ::LEU2, spo11-Y135F-HA-URA3, ndt80Δ::TRP1, RNR4-13myc::HIS3MX6 | SK1 | 5c, 5d, 5e, S9B, S9C |
| H119 | MATa, ho::LYS2, lys2, ura3,<br>leu2::hisG,<br>MATalpha, ho::LYS2, lys2, ura3,<br>leu2::hisG,<br>his4B::LEU2, arg4-Bgl<br>II<br>his4X::LEU2 (Bam)-URA3, arg4-Nsp | SK1 | S5 |
| H10963 | MATa, ho::LYS2, lys2, ura3,<br>leu2::hisG, his3::hisG, trp1::hisG,<br>hrr25-2a<br>MATalpha, ho::LYS2, lys2, URA3,<br>LEU2, his3::hisG, trp1::hisG, hrr25-2a<br>2a<br>Serine438Alanine<br>Threonine453Alanine | SK1 | S5 |
| H11031 | MATa, ho::LYS2, lys2, ura3,<br>leu2::hisG, his3::hisG, trp1::hisG<br>leu2::pURA3-TetR-GFP::LEU2,<br>HRR25-13MYC::HIS3MX6<br>MATalpha, ho::LYS2, lys2, ura3,<br>leu2::hisG, his3::hisG, trp1::hisG<br>leu2::pURA3-TetR-GFP::LEU2,<br>ura3::TETOx224::URA3, HRR25-13MYC::HIS3MX6 | SK1 | S5 |
| H7797 | MATa, ho::LYS2, lys2, ura3,<br>leu2::hisG, his3::hisG, trp1::hisG<br>MATalpha, ho::LYS2, lys2, URA3,<br>LEU2, HIS3, TRP1 | SK1 | S9A |
| H11368 | MATa, ho::LYS2, ura3, leu2::hisG,<br>his3::hisG, trp1::hisG,<br>MATalpha, ho::LYS2, lys2,<br>leu2::hisG, HIS, URA3, TRP1,<br>pph3Δ::LEU2<br>pph3Δ::LEU2 | SK1 | S9A |
| H11505 | MATa, ho::LYS2, lys2, URA3, LEU2,<br>his3::hisG, trp1::hisG<br>MATalpha, ho::LYS2, lys2, ura3,<br>leu2::hisG, HIS3, TRP1<br>pCLB2-DBP2::KanMX6<br>pCLB2-DBP2::KanMX6 | SK1 | S9A |
| H11506 | MATa, ho::LYS2, lys2, leu2::hisG,<br>HIS3, TRP1, ura3,<br>MATalpha, ho::LYS2, lys2,<br>leu2::hisG, his3::hisG, trp1::hisG,<br>URA3,<br>pph3Δ::LEU2, pCLB2-DBP2::KanMX6<br>pph3Δ::LEU2, pCLB2-DBP2::KanMX6 | SK1 | S9A |

**Supplemental Table 2:** m/z windows used for DIA

| Start | End | Width |
| --- | --- | --- |
| 350 | 385 | 35 |
| 384 | 412 | 28 |
| 411 | 434 | 23 |
| 613 | 636 | 23 |
| 454 | 474 | 20 |
| 493 | 513 | 20 |
| 552 | 572 | 20 |
| 473 | 494 | 21 |
| 512 | 533 | 21 |
| 532 | 553 | 21 |
| 433 | 455 | 22 |
| 571 | 593 | 22 |
| 592 | 614 | 22 |
| 635 | 659 | 24 |
| 658 | 683 | 25 |
| 682 | 709 | 27 |
| 708 | 738 | 30 |
| 737 | 771 | 34 |
| 770 | 808 | 38 |
| 807 | 848 | 41 |
| 847 | 899 | 52 |
| 898 | 966 | 68 |
| 965 | 1,068 | 103 |
| 1,067 | 1,650 | 583 |

**Supplemental Table 3:** DNA break-dependent phosphorylation events

| **Protein** | **Site** | **Approach for categorization** |
| --- | --- | --- |
| ABD1 | 12 | presence/absence |
| ABF1 | 189 | fold enrichment |
| ABF1 | 193 | fold enrichment |
| ABF1 | 467 | fold enrichment |
| AHP1 | 2 | presence/absence |
| AIM21 | 85 | fold enrichment |
| AKL1 | 12 | presence/absence |
| APC1 | 310 | presence/absence |
| APN1 | 346 | presence/absence |
| ASF1 | 270 | fold enrichment |
| ASF1 | 265 | presence/absence |
| ASG1 | 166 | presence/absence |
| ATG13 | 429 | presence/absence |
| ATG13 | 649 | presence/absence |
| AZF1 | 312 | presence/absence |
| AZF1 | 313 | presence/absence |
| AZF1 | 325 | presence/absence |
| BBC1 | 895 | fold enrichment |
| BBC1 | 894 | fold enrichment |
| BDF1 | 630 | fold enrichment |
| BDF1 | 626 | presence/absence |
| BIR1 | 747 | fold enrichment |
| BIR1 | 381 | fold enrichment |
| BIR1 | 751 | fold enrichment |
| BOI1 | 655 | presence/absence |
| BUB1 | 384 | presence/absence |
| BUD14 | 640 | presence/absence |
| BUD14 | 641 | presence/absence |
| BUD14 | 642 | presence/absence |
| BUD21 | 137 | fold enrichment |
| BYE1 | 210 | fold enrichment |
| CBF1 | 45 | fold enrichment |
| CBF1 | 48 | fold enrichment |
| CDC4 | 53 | presence/absence |
| CHS3 | 538 | fold enrichment |
| CIN5 | 196 | fold enrichment |
| CIN5 | 85 | presence/absence |
| CIN5 | 89 | presence/absence |
| CIN8 | 259 | fold enrichment |
| CIN8 | 261 | fold enrichment |
| CIT2 | 21 | presence/absence |
| CKB2 | 12 | presence/absence |
| CMR1 | 224 | fold enrichment |
| CMR1 | 79 | fold enrichment |
| CUS1 | 104 | presence/absence |
| CYR1 | 241 | presence/absence |
| DAL81 | 17 | fold enrichment |
| DBF4 | 84 | fold enrichment |
| DBF4 | 473 | fold enrichment |
| DBF4 | 496 | presence/absence |
| DBF4 | 501 | presence/absence |
| DBP7 | 56 | presence/absence |
| DCP2 | 686 | fold enrichment |
| DDC1 | 558 | presence/absence |
| DGK1 | 3 | presence/absence |
| DIF1 | 102 | presence/absence |
| DIF1 | 104 | presence/absence |
| DMA2 | 206 | presence/absence |
| DPB4 | 183 | fold enrichment |
| DPS1 | 301 | presence/absence |
| DST1 | 115 | fold enrichment |
| DUN1 | 10 | presence/absence |
| ECM11 | 150 | fold enrichment |
| ELC1 | 2 | presence/absence |
| ENT1 | 160 | fold enrichment |
| ENT1 | 163 | fold enrichment |
| ENT1 | 392 | presence/absence |
| ENT2 | 468 | presence/absence |
| ENT2 | 470 | presence/absence |
| ESL1 | 174 | presence/absence |
| EXO1 | 663 | presence/absence |
| EXO1 | 664 | presence/absence |
| FBP1 | 13 | fold enrichment |
| FIN1 | 68 | fold enrichment |
| GAL11 | 789 | presence/absence |
| GCD1 | 296 | presence/absence |
| GCD14 | 302 | fold enrichment |
| GCD14 | 299 | presence/absence |
| GFA1 | 332 | presence/absence |
| GIP2 | 117 | presence/absence |
| GIP2 | 120 | presence/absence |
| GLC7 | 3 | fold enrichment |
| GPA2 | 23 | presence/absence |
| GTT1 | 56 | fold enrichment |
| HAS1 | 12 | fold enrichment |
| HAS1 | 14 | presence/absence |
| HBT1 | 1034 | fold enrichment |
| HBT1 | 1036 | fold enrichment |
| HBT1 | 965 | fold enrichment |
| HED1 | 40 | presence/absence |
| HED1 | 42 | presence/absence |
| HHT1 | 58 | presence/absence |
| HOG1 | 176 | fold enrichment |
| HOP1 | 22 | presence/absence |
| HPC2 | 307 | fold enrichment |
| HPC2 | 388 | presence/absence |
| HRB1 | 27 | fold enrichment |
| HRR25 | 438 | presence/absence |
| HRT1 | 15 | presence/absence |
| HSP12 | 21 | fold enrichment |
| HSP12 | 24 | fold enrichment |
| HSP12 | 59 | fold enrichment |
| HSP26 | 208 | presence/absence |
| HSP26 | 211 | presence/absence |
| HTA1 | 129 | fold enrichment |
| HXT2 | 17 | fold enrichment |
| IGD1 | 64 | fold enrichment |
| IGD1 | 83 | presence/absence |
| IGO2 | 119 | fold enrichment |
| INO1 | 2 | presence/absence |
| IOC2 | 629 | presence/absence |
| IPL1 | 5 | fold enrichment |
| IRR1 | 14 | fold enrichment |
| IRR1 | 12 | fold enrichment |
| IRR1 | 21 | presence/absence |
| IVY1 | 32 | presence/absence |
| LEO1 | 339 | presence/absence |
| MAF1 | 179 | presence/absence |
| MAK21 | 2 | presence/absence |
| MBP1 | 133 | fold enrichment |
| MCM10 | 66 | presence/absence |
| MCM4 | 52 | fold enrichment |
| MDS3 | 621 | fold enrichment |
| MEC3 | 368 | fold enrichment |
| MEC3 | 369 | fold enrichment |
| MEK1 | 486 | presence/absence |
| MEK1 | 356 | presence/absence |
| MET12 | 120 | presence/absence |
| MLH1 | 441 | fold enrichment |
| MMS4 | 2 | fold enrichment |
| MON1 | 130 | fold enrichment |
| MOT1 | 679 | presence/absence |
| MRC1 | 121 | presence/absence |
| MRE11 | 558 | fold enrichment |
| MSC3 | 57 | fold enrichment |
| MSC3 | 64 | fold enrichment |
| MSH6 | 102 | fold enrichment |
| MTL1 | 481 | fold enrichment |
| MTR4 | 84 | fold enrichment |
| NAB2 | 2 | presence/absence |
| NET1 | 1025 | fold enrichment |
| NET1 | 1026 | fold enrichment |
| NET1 | 1024 | fold enrichment |
| NET1 | 440 | presence/absence |
| NET1 | 439 | presence/absence |
| NGG1 | 231 | presence/absence |
| NNK1 | 689 | fold enrichment |
| NOG2 | 60 | fold enrichment |
| NOT3 | 446 | presence/absence |
| NPL3 | 224 | fold enrichment |
| NUP53 | 95 | fold enrichment |
| NUP60 | 10 | fold enrichment |
| NUP60 | 458 | fold enrichment |
| NUP60 | 460 | fold enrichment |
| NUP60 | 483 | presence/absence |
| NUR1 | 441 | presence/absence |
| OLA1 | 67 | presence/absence |
| PAP1 | 550 | fold enrichment |
| PCT1 | 67 | fold enrichment |
| PEX3 | 118 | fold enrichment |
| PEX3 | 119 | fold enrichment |
| PFK1 | 3 | fold enrichment |
| PIL1 | 45 | fold enrichment |
| PRI2 | 16 | fold enrichment |
| PRI2 | 17 | fold enrichment |
| PUP2 | 128 | fold enrichment |
| RAD1 | 40 | fold enrichment |
| RAD14 | 360 | fold enrichment |
| RAD16 | 109 | presence/absence |
| RAD17 | 350 | presence/absence |
| RAD23 | 121 | presence/absence |
| RAD24 | 637 | fold enrichment |
| RAD26 | 27 | presence/absence |
| RAD52 | 205 | presence/absence |
| RAD54 | 132 | presence/absence |
| RAD7 | 64 | fold enrichment |
| RCK2 | 187 | presence/absence |
| RDI1 | 9 | presence/absence |
| REC114 | 55 | fold enrichment |
| RED1 | 483 | fold enrichment |
| RED1 | 484 | fold enrichment |
| RED1 | 486 | fold enrichment |
| RED1 | 473 | fold enrichment |
| RED1 | 517 | fold enrichment |
| RED1 | 518 | fold enrichment |
| RFA2 | 115 | presence/absence |
| RFM1 | 83 | presence/absence |
| RFX1 | 226 | presence/absence |
| RFX1 | 173 | presence/absence |
| RGT1 | 284 | fold enrichment |
| RIF1 | 1362 | fold enrichment |
| RIM15 | 555 | presence/absence |
| RPC53 | 234 | presence/absence |
| RPS16A | 15 | fold enrichment |
| RPS2 | 30 | fold enrichment |
| RPS7B | 14 | presence/absence |
| RRB1 | 5 | fold enrichment |
| RRM3 | 125 | presence/absence |
| RRP12 | 1050 | presence/absence |
| RRP36 | 41 | fold enrichment |
| RRP36 | 42 | fold enrichment |
| RSC9 | 44 | presence/absence |
| RTF1 | 17 | presence/absence |
| RTF1 | 2 | presence/absence |
| RTG3 | 241 | fold enrichment |
| RTG3 | 246 | fold enrichment |
| RTG3 | 269 | presence/absence |
| RTG3 | 236 | presence/absence |
| RTS2 | 160 | presence/absence |
| RTT107 | 591 | fold enrichment |
| RTT107 | 593 | fold enrichment |
| RTT107 | 735 | fold enrichment |
| RTT107 | 800 | fold enrichment |
| RTT107 | 806 | fold enrichment |
| RTT107 | 255 | presence/absence |
| SEC16 | 2139 | presence/absence |
| SEC16 | 2141 | presence/absence |
| SEC3 | 254 | presence/absence |
| SEC3 | 256 | presence/absence |
| SEF1 | 273 | presence/absence |
| SEG1 | 48 | presence/absence |
| SFT2 | 2 | presence/absence |
| SGF73 | 19 | fold enrichment |
| SGF73 | 22 | presence/absence |
| SGO1 | 421 | fold enrichment |
| SGO1 | 423 | fold enrichment |
| SGO1 | 426 | fold enrichment |
| SGS1 | 482 | presence/absence |
| SGS1 | 606 | presence/absence |
| SGV1 | 417 | fold enrichment |
| SHP1 | 226 | presence/absence |
| SIR2 | 23 | presence/absence |
| SIR3 | 263 | presence/absence |
| SIR4 | 692 | fold enrichment |
| SIR4 | 342 | presence/absence |
| SKI7 | 88 | fold enrichment |
| SKI7 | 90 | fold enrichment |
| SLD2 | 150 | fold enrichment |
| SLD2 | 151 | fold enrichment |
| SLI15 | 97 | presence/absence |
| SLK19 | 216 | fold enrichment |
| SLM1 | 157 | fold enrichment |
| SLM1 | 158 | fold enrichment |
| SMB1 | 67 | fold enrichment |
| SPC105 | 144 | presence/absence |
| SPC110 | 60 | fold enrichment |
| SPC24 | 2 | presence/absence |
| SPC29 | 230 | fold enrichment |
| SPC29 | 231 | fold enrichment |
| SPC29 | 248 | fold enrichment |
| SPC29 | 249 | fold enrichment |
| SPC29 | 250 | fold enrichment |
| SPG4 | 93 | fold enrichment |
| SPO13 | 139 | fold enrichment |
| SPO13 | 134 | presence/absence |
| SPP1 | 18 | fold enrichment |
| SPP41 | 563 | fold enrichment |
| SPP41 | 564 | fold enrichment |
| SPT7 | 78 | fold enrichment |
| SRP40 | 394 | presence/absence |
| SSD1 | 489 | presence/absence |
| SSN2 | 748 | presence/absence |
| SSN2 | 521 | presence/absence |
| STB3 | 337 | fold enrichment |
| STB3 | 2 | presence/absence |
| STE20 | 192 | presence/absence |
| STE20 | 195 | presence/absence |
| STU1 | 1000 | fold enrichment |
| SUB2 | 2 | fold enrichment |
| SUB2 | 12 | fold enrichment |
| SUB2 | 13 | fold enrichment |
| SUM1 | 722 | fold enrichment |
| SUM1 | 858 | fold enrichment |
| SUM1 | 859 | fold enrichment |
| SWI3 | 88 | fold enrichment |
| SWI3 | 185 | fold enrichment |
| SYH1 | 687 | fold enrichment |
| TAF3 | 345 | fold enrichment |
| TAF3 | 346 | fold enrichment |
| TAF8 | 212 | fold enrichment |
| TAF8 | 215 | fold enrichment |
| TCB3 | 1340 | fold enrichment |
| THO1 | 72 | fold enrichment |
| THO1 | 68 | presence/absence |
| THS1 | 2 | presence/absence |
| TIF4632 | 913 | fold enrichment |
| TIF5 | 184 | presence/absence |
| TOA2 | 102 | presence/absence |
| TOA2 | 95 | presence/absence |
| TOP1 | 24 | fold enrichment |
| TPO3 | 52 | presence/absence |
| TRM10 | 16 | presence/absence |
| TSL1 | 157 | fold enrichment |
| UBC5 | 12 | fold enrichment |
| UBP1 | 776 | fold enrichment |
| UBP13 | 463 | fold enrichment |
| UGX2 | 186 | fold enrichment |
| ULP1 | 264 | fold enrichment |
| ULP1 | 48 | fold enrichment |
| ULP2 | 773 | fold enrichment |
| ULP2 | 929 | fold enrichment |
| ULP2 | 936 | fold enrichment |
| ULP2 | 937 | fold enrichment |
| ULS1 | 204 | fold enrichment |
| UNG1 | 23 | presence/absence |
| USV1 | 369 | presence/absence |
| UTP22 | 10 | fold enrichment |
| UTP5 | 629 | presence/absence |
| UTP5 | 637 | presence/absence |
| VTC2 | 187 | presence/absence |
| XRN1 | 1510 | fold enrichment |
| XRS2 | 783 | presence/absence |
| YBR285W | 117 | presence/absence |
| YCS4 | 495 | presence/absence |
| YDL199C | 61 | fold enrichment |
| YDR090C | 306 | presence/absence |
| YDR090C | 309 | presence/absence |
| YDR239C | 74 | presence/absence |
| YDR239C | 681 | presence/absence |
| YER079W | 10 | presence/absence |
| YER079W | 9 | presence/absence |
| YET1 | 172 | presence/absence |
| YGR130C | 803 | fold enrichment |
| YGR130C | 44 | presence/absence |
| YHR097C | 69 | fold enrichment |
| YJL206C | 90 | presence/absence |
| YLR257W | 238 | presence/absence |
| YPQ2 | 136 | presence/absence |
| ZIP1 | 549 | presence/absence |
| ZIP1 | 75 | presence/absence |
| ZIP1 | 546 | presence/absence |
| ZIP1 | 865 | presence/absence |

**Supplemental Table 4:** Phosphosites enriched in *spo11-YF*

| **Protein** | **Site** | **Approach for categorization** |
| --- | --- | --- |
| ABF1 | 159 | presence/absence |
| ACF4 | 9 | presence/absence |
| CRZ1 | 453 | presence/absence |
| CUS1 | 114 | presence/absence |
| DNA2 | 17 | fold enrichment |
| DPS1 | 30 | presence/absence |
| IMH1 | 25 | presence/absence |
| KEL1 | 1019 | presence/absence |
| KKQ8 | 113 | presence/absence |
| LIF1 | 261 | presence/absence |
| MLF3 | 145 | presence/absence |
| PDS1 | 212 | presence/absence |
| PDS1 | 213 | presence/absence |
| RPL24A;RPL24B | 83;83 | presence/absence |
| RPL24A;RPL24B | 86;86 | presence/absence |
| SEC3 | 301 | presence/absence |
| THO1 | 48 | presence/absence |
| UFD1 | 314 | presence/absence |
| VPS27 | 280 | presence/absence |
| YFR016C | 49 | presence/absence |
| YJL163C | 59 | presence/absence |
| YJL163C | 61 | presence/absence |
| YOL036W | 547 | presence/absence |
| YPL150W | 900 | presence/absence |
| ARG82 | 98 | fold enrichment |
| CKI1 | 54 | fold enrichment |
| GET2 | 60 | fold enrichment |
| PIN4 | 638 | fold enrichment |
| ARO80 | 96 | fold enrichment |
| DNA2 | 236 | fold enrichment |
| DNA2 | 237 | fold enrichment |
| ESC2 | 71 | fold enrichment |
| ESC2 | 73 | fold enrichment |
| ESC2 | 76 | fold enrichment |
| PIB2 | 68 | fold enrichment |
| PIB2 | 70 | fold enrichment |
| REC104 | 110 | fold enrichment |
| RHO4 | 54 | fold enrichment |
| RHO4 | 55 | fold enrichment |
| RTT107 | 720 | fold enrichment |
| STU1 | 1113 | fold enrichment |
| UBP1 | 652 | fold enrichment |
| XRS2 | 349 | fold enrichment |
| AVO1 | 144 | presence/absence |
| BNR1 | 621 | presence/absence |
| BUL1 | 195 | presence/absence |
| DRE2 | 212 | presence/absence |
| EDE1 | 1062 | presence/absence |
| FAS2 | 1440 | presence/absence |
| GYP6 | 436 | presence/absence |
| HOP1 | 553 | presence/absence |
| MCM10 | 18 | presence/absence |
| MCM10 | 68 | presence/absence |
| OSH3 | 447 | presence/absence |
| OSH3 | 449 | presence/absence |
| OSH3 | 452 | presence/absence |
| PKH1 | 303 | presence/absence |
| PXA2 | 776 | presence/absence |
| RAD7 | 118 | presence/absence |
| REC107 | 37 | presence/absence |
| RGA2 | 733 | presence/absence |
| RGC1 | 1033 | presence/absence |
| RHO4 | 58 | presence/absence |
| RRP9 | 76 | presence/absence |
| SEC16 | 1633 | presence/absence |
| SEC16 | 1634 | presence/absence |
| SEC16 | 1640 | presence/absence |
| SGS1 | 643 | presence/absence |
| SGT1 | 166 | presence/absence |
| SKG3 | 814 | presence/absence |
| SUI2 | 52 | presence/absence |
| SVL3 | 551 | presence/absence |
| SWI3 | 230 | presence/absence |
| SWI3 | 231 | presence/absence |
| SWI3 | 91 | presence/absence |
| TSL1 | 137 | presence/absence |
| VHS3 | 264 | presence/absence |
| VID27 | 417 | presence/absence |
| YAT2 | 848 | presence/absence |
| YEL043W | 895 | presence/absence |
| YPL150W | 807 | presence/absence |

